# SeQuoIA: a single-cell BCR sequencing analysis pipeline for tracking selection mechanisms in germinal centres

**DOI:** 10.1101/2025.02.13.638132

**Authors:** Eglantine Hector, Pierre Milpied

**Author notes:** Corresponding authors: Eglantine Hector, Pierre Milpied.

## Abstract

Germinal centres (GCs) are specialized structures where B cells undergo iterative steps of B-cell receptor (BCR) somatic hypermutation and selection of best antigen binders in a darwinian-like fashion. The accelerated evolutionary process leads to the production of high-affinity antibodies that are crucial for robust and long-term humoral immunity. Within this frame, single-cell BCR sequencing analysis is a method of choice to track GC B cell dynamics as somatic mutations can be utilised as an *in vivo* molecular tracer. Herein, we present SeQuoIA, a start-to-finish pipeline for the analysis of BCR repertoire sequencing data at the single-cell level, including improved clonotype assignment and phylogeny reconstruction. Most importantly, we introduce a new method for the inference of BCR-driven selection pressure based on somatic mutation patterns, that was validated with biological data. With this pipeline, we explored public datasets and proposed new selection mechanisms in GCs.

**Significance:** Our pipeline should contribute to a better understanding of the basic biology of GC dynamics, and potentially help in laboratory animal usage reduction. Clinical applications could include assessment of vaccine efficacy, monitoring of B cell anti-tumoral responses, and identification of BCR-mediated processes in B cell lymphomas.

## Introduction

B cells are major actors of the adaptive immune response against pathogens. These cells are armed with immunoglobulins (Ig), in the form of receptor (BCR) or secreted antibodies, which can bind foreign antigens with high affinity and specificity. These features are acquired through successive diversification and selection steps in the bone marrow (BM) and secondary lymphoid organs (SLO). VDJ recombination during early B cell development generates combinatorial diversity of BCR repertoires. Upon infection or immunisation, circulating mature B cells can undergo additional development steps in specialised micro-anatomic niches called germinal centres (GCs). GC entry is determined by B cell ability to bind the antigen (Ag) and receive T cell signals. This bottleneck provides a first clonal selection step. Within GCs, B cells migrate between two compartments in the cyclic process of affinity maturation (AM). They diversify their BCR through somatic hypermutation (SHM) in the dark zone (DZ). Point mutations are introduced by activation cytidine deaminase (AID) enzyme. The binding properties of newly generated receptors are tested in the light zone (LZ), where B cells compete for antigen access and those harbouring adequate BCR affinity are selected in a darwinian-like process. After several rounds of diversification and selection, B cells may eventually exit GCs and differentiate into memory B cells (MBCs) or plasma cells (PCs).

How Ag binding properties translate into increased fitness is not fully understood. B cell selection has been mostly scrutinised through the lens of antigen presentation to follicular helper T cells (T_FH_), which provide cues (CD40L, IL4, IL21) involved in B cell proliferation and differentiation. Within this framework, GC B cells failing to receive T_FH_ cues are assumed to die by neglect^1^. The role of other signals is yet to be fully explored. Fate decisions (differentiation, death and recycling) and the signals that govern them is still an area of active research. In most experimental model systems, it has been shown that antibody-producing PC differentiated preferentially from GC B cells expressing BCR with high affinity for the antigen, although in polyclonal responses to complex antigens broad ranges of BCR affinities may end up in the GC-derived PC compartment^2,3^. Within this context, the extent to which PC differentiation relies on BCR affinity has not been settled.

Single cell datasets including gene expression and Ig sequence data constitute an insightful approach for the exploration of GC reaction (GCR) selection mechanisms, as they link clonal and mutation information to a transcriptional state characteristic of cell subset or recent signal transduction. Many tools have already been developed for processing Ig sequencing (IgSeq) datasets, identifying clonal families and tracing Ig sequence evolution through phylogenetic tree reconstruction. For an exhaustive list of such tools as well as typical repertoire analyses, the reader is referred to several reviews^4,5,6^. Despite the large amount of softwares dedicated to IgSeq analysis, few attempts have been made to infer BCR-driven selection pressure from SHM patterns^7^. Tools integrating gene expression, Ig heavy and light chain information are lacking, although high-throughput integrative single cell sequencing made strides over the past few years.

We developed a bioinformatic pipeline for comprehensive BCR repertoire analysis at the single cell level: **Se**lection **Qu**antification in **I**ntegrative **A**IRR data (SeQuoIA). SeQuoIA achieved improved clonotype assignment and phylogenetic reconstruction accuracy over state-of-the-art existing tools. Most importantly, *SeQuoIA* uniquely leverages BCR clonal tree knowledge to quantify selection based on Ig mutation patterns. Finally, our pipeline links BCR features to transcriptional states. In this article, we applied SeQuoIA to publicly available datasets and we show that the light chain is a valuable source of information, both for phylogeny reconstruction and tracking selection. We also demonstrate that SeQuoIA is a powerful predictive tool to generate new hypotheses on BCR-driven selection mechanisms, in the LZ but also subsequently in the DZ cells or after commitment to PC differentiation.

## Material and Methods

### SeQuoIA workflow

The complete pipeline is summarised in Fig.1 and pipeline code description can be found in Supp. Table 1.

**Figure 1.**
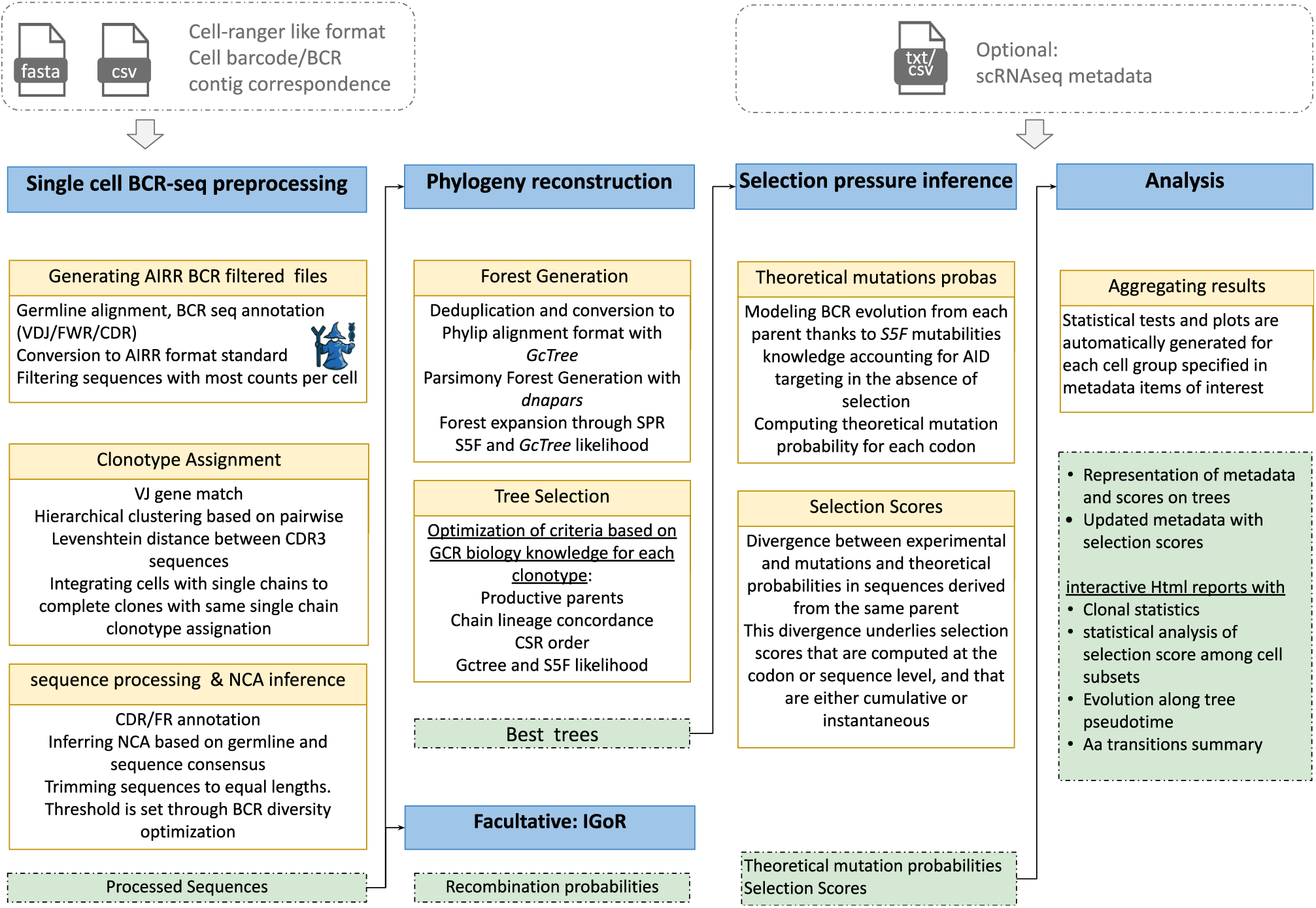
SeQuoIA pipeline. Inputs and outputs are depicted in gray and green respectively. Each yellow box corresponds to scripts described in supplementary tables.

#### Sequencing data pre-processing and annotation

Sequencing error correction, read assembly and read quality filtering are expected to be done by the user prior to SeQuoIA. The SeQuoIA pipeline takes as input assembled contigs in fasta format and an annotation file linking Ig contig information to cell barcodes. These inputs are typically generated with CellRanger V(D)J but could easily be obtained with other BCR sequencing data preprocessing methods. Additional but highly recommended input includes cell information from GEX data analysis (e.g. metadata from Seurat object). Input formatting requirements are detailed in Supp. Table 2.

Contigs are aligned onto a reference genome (mouse or human) and annotated with the *Immcantation* suite^8^, including V, D, J gene segment usage and functional annotation (framework regions FRW and complement determining regions CDR). Contig filtering is based on alignment quality (non-missing VJ portions) and cellular data (contig with higher UMI counts per chain is kept). All intermediate outputs are in Adaptive Immune Receptor Repertoire (AIRR) format^9^.

#### Clonotype assignation

Clonally related sequences descending from the same B cell are identified with classical similarity approaches. Similarity based grouping on sequences sharing the same V and J allele is performed for each chain with a hierarchical clustering on CDR3 aa sequences hamming distance matrix, and a 80% similarity cutoff is applied. The added value of SeQuoIA as compared to most clonal assignment tools is the pairing of heavy and light chain information through single-cell ID (10x barcode or well position). Heavy and light chain clones are combined into single consistent clones. Cells missing an Ig chain can be incorporated into clonal groups defined with both chain information if the corresponding chain meets the above-mentioned criteria. Sequences with indels are usually filtered out from clonal families at this stage as frameshifts lead to dramatic changes in CDR3 aa sequence. Clonotype assignment can be performed at the dataset level or for each donor independently, if donor information is provided by the user. The donor-restricted approach allows computation time to be reduced and increases the likelihood that clonal cells are indeed related, even in the case of “public” clones or murine genetically engineered B cells harbouring the same V(D)J combination.

#### Ancestral sequence reconstruction

For each clonotype, a clone ancestor is inferred. This ancestor corresponds either to the germline (*e.g.* naive B cell derived clones) or the Nearest Common Ancestor (NCA), a virtual or sampled sequence from which all sampled sequences should derive. The NCA is relevant in memory B cell derived clones or clonal families that are advanced in the affinity maturation process.

At this stage, germline sequences lack junction information, which cannot be inferred from genome alignment (*cf.* pre-processing section). Junction sequences are thus filled with a 50% consensus of least mutated contigs (first two ranks).

NCA reconstruction is based on a strict consensus of least mutated sequences for each nucleotide. Germline nucleotides are taken in positions lacking consensus. Ancestor sequence productivity is ensured by SeQuoIA.

#### Phylogeny reconstruction

Ancestral relationships within clonal families are inferred with a maximum parsimony based phylogenetic reconstruction of heavy and light chains. Trees are rooted either to the NCA or the germline (if the distance to the germline in non-junction sequence is inferior to 3). Identical Ig sequences are made non redundant with--*deduplicate* utility from *GcTree*^10^ (v4.0.0), which also summarises the information of unique Ig sequences abundance. Phylip *dnapars* is called to generate a set of most parsimonious trees, henceforth termed the “Parsimony Forest”. Non parsimonious trees are derived from the parsimonious ones through a prune and regraft (SPR) approach where nodes can be moved upstream of the trees. One pair of heavy and light chain trees is selected within the extended forest (featuring parsimonious and SPR-derived trees). Tree selection is informed by several criteria based on current knowledge of affinity maturation biology which are, by order of priority:

- <u>Mutation load and reversions:</u> reverse mutations are unexpected except in pathological generation of bystander and autoreactive cells as evidenced in BCR sequencing at different timepoints in WT and ALPS mouse models^11^. More generally, trees with least mutation ‘aberrations’ (reversion, decreased mutation load along tree progression axis) are pre-selected with SeQuoIA.
- <u>Parent productivity:</u> DZ B cells were shown to have a quick BCR turnover and to undergo apoptosis before reaching the LZ if a deleterious mutation led to a destabilisation of the BCR complex^12^. As a result, cells expressing non-productive sequences should not be able to have a descendance. SeQuoIA takes into account the need for tonic BCR signalling to persist in GCs by filtering out trees with non-productive (*i.e.* with a stop codon or a frameshift) internal nodes, whether they are observed or inferred.
- <u>CSR parsimony</u>: this criterion takes into consideration the isotypes of observed nodes and minimises the number of independent CSR events along the tree. CSR parsimony is already implemented in the *GcTree* package (*isotype_parsimony* function).
- <u>Chain concordance</u>: this criterion is based on paired heavy and light chain data recovered in scRNAseq. Heavy and light chain sequences expressed within the same cell should have the same lineage in both heavy and light chain trees. However, checking the concordance of lineages is not trivial due to an unbalanced evolution speed between both chains. As a result, heavy chains tend to have many divergent sequences matched to the same light chain sequence. SeQuoIA traces back each node with paired data in heavy and light chain trees and checks whether:

- nodes have shared parents in both trees. If not, unshared parent could correspond to sequences derived from the same grandparent where heavy and light chains mutations were non concomitant. SeQuoIA thus extends the search to grandparents in heavy and light chain trees. Otherwise, the chain pairs are considered as not directly related in heavy and light chain trees. The tree pair combination is excluded from the exploration space for subsequent filtering criteria.
- Paired sequences are in the same order from ancestor to terminal leaves. If the order is not conserved, the tree pair is excluded from the exploration space.

SeQuoIA then ranks the remaining trees so as to choose the tree pairs that maximise the overlap in chain combination lineages. The degree of overlap is quantified with a modified jaccard index normalised by the total number of tree pairs:

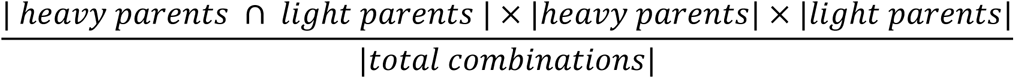

The chain concordance-based selection process is illustrated in Supp. Fig.1.

- <u>CSR order</u>: CSR is an irreversible genomic rearrangement involving the excision of exons encoding alternative Ig classes. CSR proceeds from upstream to downstream exons along the IGH constant region locus in the following order: IgD, IgM, IgG3, IgG1, IgG2b, IgG2c, IgE, IgA in mouse, and IgD, IgM, IgG3, IgG1, IgA1, IgG2, IgG4, IgE, IgA2 in human. The number of breaches from this order is counted by SeQuoIA and used to rank trees that passed previous filters. Since most CSR are thought to occur prior to GC entry ^13^, these CSR events should not occur during affinity maturation. However, they may occur during precursor activation and possibly during differentiation into effector B cells.
- <u>GcTree likelihood</u>: the *GcTree* tool incorporates node abundance (*i.e.* identical Ig sequence frequencies) to rank trees in terms of likelihood within the parsimony forest. This criterion is based on the premise that the more an Ig sequence is represented, the more likely it is to have a descendance. Most expanded nodes should thus be located upstream of the tree axis.
- <u>Mutability Parsimony</u>: Equally parsimonious trees could also be ranked according to the likelihood of SHM mutability patterns, as implemented in SAMM^14^ and later incorporated in the *GcTree* pipeline. However, clonotypes subject to selection should deviate from the basal AID mutability patterns. This criterion is thus used as the last filter.

#### Mutation Simulation

Parent sequences (i.e. tree internal nodes) are used to initialise several independent models recapitulating point mutation accumulation in the absence of selection. Inferred internal nodes can be taken if the selected tree has a non-ambiguous topology. Mutations are introduced in parent sequences until the mutation level of its daughters is reached, and taking into account SHM intrinsic bias which has been characterised for hundreds of 5-mer nucleotide patterns^15^. In practice, these mutations are applied with the shmulateSeq function of the *SHazaM* package, with either previously published human targeting model^15^ or our own mouse targeting model depending on the species configuration parameter. SeQuoIA applies mutations sequentially to update mutability patterns and ensures sequence productivity at each step. Newly *in silico* generated sequences are translated to check whether the mutation is synonymous or not. The mutation steps are repeated in 2000 replicates and mutation indices are gathered to estimate conditional probabilities of a site being mutated:

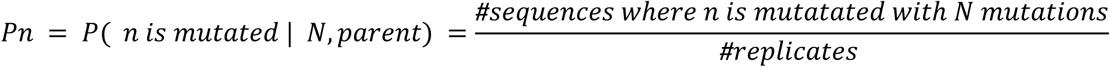

where n is the nucleotide index, *parent* is the parental sequence used for initialization, and *N* the total (synonymous and non-synonymous) number of mutations accumulated from this parent. *N* is low in trees where intermediate states could be captured. In the default mode, the pipeline does not assume that only non-synonymous (NS) mutations affect BCR fitness and all mutations are considered. SeQuoIA generates figures as QCs to compare *Pn* in well described SHM hotspot (WRC) and coldspot (SYC) (Supp. Fig.2).

#### Selection Quantification

Experimental sequences mutation analysis includes:

- Mutation load: since parent and ancestor
- Mutation localization: nucleotide or codon index in RNA or protein sequences respectively
- Mutation effect: synonymous or non-synonymous

Theoretical mutation probabilities estimated in the previous step are used to compute selection scores. A codon selection score summarises the degree of expectancy of mutation or non-mutation in mutated and conserved codons respectively:

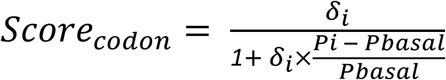

where

- δ_*i*_ = 1 if codon i is affected by a NS mutation, δ_*i*_ = 0 otherwise
- *P_i_* the probability that at least one of the nucleotides in codon i is non-synonymously mutated. *P_i_* =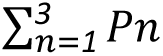
- *Pbasal* is the mean over all Pi within the sequence. This normalisation factor allows to visualise the sites of appreciable positive and negative selection along a BCR sequence in a single plot.

Intuitively, the presence of a mutation although it was unlikely to occur according to SHM model (*P_i_*> Pbasal) may be an indicator of positive selection (Score_codon_ > 1). Conversely, unmutated sites in mutation hotspot may be an indicator of negative selection ((Score_codon_ <-1).

We use Kulback-Leibler (KL) divergence between observed mutations and the theoretical mutabilities to estimate positive selection imprint on the whole Ig sequence. The divergence of observed mutation from the model is normalised by the entropy to compare sequences with different ancestries and mutation loads. Sequence scores account either for instantaneous or cumulated selection imprints thanks to the phylogeny knowledge.

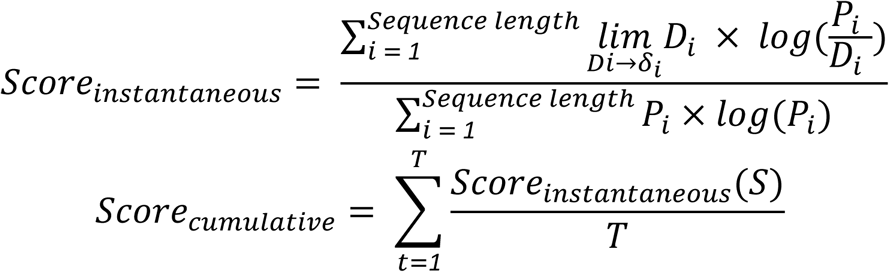

with *T* the number of tree layers until node S.

By default, unless stated otherwise, selection scores will refer to the instantaneous selection score in this manuscript.

#### Result aggregation and analysis

Experimental and transcriptomic data is incorporated into the analysis to compare selection between groups of cells or along continuous variables (tree pseudo-time, distance to germline and clone size are analysed by default). Details on generated outputs and script details can be found in Supp. Table 1. This report only provides basic analyses. The user is strongly encouraged to check tree quality (as in Supp. Fig.3) and perform gene expression UMI-based filtering prior to custom analysis.

#### Interface with IGOr

SeQuoIA can incorporate IGOr^16^ to estimate VDJ combination probabilities of each clonotype. Examples of applications are provided in Supp. Fig.4. Generation probabilities (Pgen) can serve as a proxy of clonal selection (as in pre-GC steps), on the basis that low probability BCRs could be enriched through selection steps in GC and the follicle. This approach was validated in Supp. Fig.4B, where Pgens were strongly predictive of experimental affinity measurements.

#### Resources, environment and reproducibility

The SeQuoIA pipeline is based on the *snakemake*^17^ workflow management system. *Snakemake* executes interdependent tasks (rules) and ensures reproducibility of the workflow by deploying dependencies stored in a compiled format. *Docker* images for each *snakemake* rule were thus converted to the *Singularity* ‘sif’ format. The SeQuoIA pipeline can be run locally or on a cluster. CPU resources to be used are customizable. Job parallelization is automated.

### Construction of AID mutability model

#### Sequencing datasets

The human model for SHM 5-mers mutability and substitution probabilities was already available through the Shazam package. Mouse model accounting for SHM targeting in the heavy chain was still lacking. To compute mutation probabilities with minimal selection interference, we chose an IgSeq dataset from Peyer’s Patches spontaneous GCs from germ-free mice (Supp. Fig.5A). Indeed, in the absence of gut microbiota antigens, selection for GC entry was shown to be severely dampened^18^.

#### Mutability and substitution matrices generation

The model for murine Ig sequences was built according to a standard workflow from the Immcantation suite. One sequence per clone was selected to avoid biases due to clonal expansions. Substitution rates were inferred with the *createSubstitutionMatrix*. 5-mer mutabilities were inferred with the *createMutabilityMatrix* function, considering all mutations (synonymous and non-synonymous). Both outputs were combined into a single model using the *createTargetingModel* function. 5-mer mutabilities are displayed in Supp. Fig.5B.

### scRNAseq processing and annotation

Public integrative datasets used in this study are detailed in Supp. Table 3. If provided in the form of a Seurat object, additional cell filterings were applied (at least 300 features, mitochondrial content < 5% of genes expressed). A standard Seurat (v5.0.3) workflow was applied, including cell cycle regression, dimensionality reduction and clustering. Clustering resolution was determined with an automated script based on cell cluster assignment stability with the *Clustree* package. Supervised cell annotation was performed with cluster marker genes as depicted in Supp. Fig.6, Supp. Fig.7 and Supp. Fig.8.

#### Specificities of human Covid dataset

Same QC filters were applied as described above. However, the donor effect was still noticeable in the original dataset (Supp. Fig.8B). Donor-based integration was performed with the semi-supervised method STACAS^19^. Original annotations (PCs, GC, Naive, Memory B cells) were used to refine anchor usage. GC B cells were pre-annotated into global LZ and DZ categories and cells with intermediate phenotypes were labelled as “unknown” and not taken into account for inconsistent anchor rejection. Histone, ribosomal, immunoglobulin and cell cycle coding genes were excluded from variable features prior to integration. Additional difficulties arose from this dataset, due to concomitant and intricate GC reaction and massive extrafollicular (EF) response. Clustering was performed with the same variable features as for the integration. Clusters were annotated based on marker genes and 3D UMAP visualization, where 3 “paths” to terminally differentiated PC clusters stood out (Supp. Fig.8C). These transitional clusters were all made of *XBP1*^+^ cells and were annotated as follow:

- <u>prePC EF</u>: emerging from the naive B cell cluster and CD38^neg^, suggesting that they are not GC-experienced. Not BLIMP1^+^ like other prePC clusters.
- <u>prePC</u>: emerging from the LZ
- <u>prePC DZ:</u> very small cluster of proliferating cells emerging from DZ recycling cells. Whether they are related to prePC remained unclear to us.

Likewise, an intermediary CD38**^-^** cluster bridged naive and GC “meta” clusters in the 3D umap and was hence annotated as “pre-GC”. Despite our efforts to refine original annotations, some clusters remained ambiguous and displayed marked heterogeneity. Among them, the MYC+ intermediate cluster between LZ and DZ B cells was dominated by CD38**^-^** and was the entry point of pre-GC cells (Supp. Fig.8D). This cluster thus likely corresponded to both rare selected LZ-to-DZ B cells and newly recruited GC entrants starting their first round of mutation within GCs. For some analysis, *bona fide* LZ-to-DZ were enriched by selecting cells that were not directly connected to the ancestor and thus should have completed at least one GC cycle. We also endeavoured to segregate EF and GC-derived PCs. However, in the absence of GC-origin lineage tracing system like the one used in the murine datasets, PC origin was not obvious. Since PC cluster sub-clustering did not seem to segregate two relevant PC categories, putative EF PCs were identified based on clonotype overlap with prePC EF cluster. Remaining PCs were labelled “PC other” and should be enriched in cells of GC origin. All analyses performed with this re-annotated dataset are gathered in Supp. Fig.11.

## Results

### SeQuoIA improves Ig phylogeny accuracy with biologically-based optimization and by reducing the occurrence of dubious clonotypes

Many tree reconstruction methods have been developed to recover Igs evolutionary dynamics. Albeit based on pioneering work in ecological sciences, most of the tools accommodate BCR biology. Indeed, as GC B cells proliferate and mutate in the short evolutive time span of GCR, both child and parent may coexist at the time of sampling, hence the incorporation of internal nodes. In particular, GcTree^10^ achieved good rankings in simulation-based benchmarking studies^20,21^ and thus served as a basis for the implementation of the SeQuoIA pipeline. This tool consists in an informed maximum parsimony method, where the best tree within the parsimony forest is picked based on node size distribution and the compliance to mutation models. As most accurate parent to daughter relationships are needed in the SeQuoIA mutation model (see next section), we added additional optimization steps to guide tree selection. Due to the presence of mutation hotspots and the high mutation rate in Ig sequences, the number of mutations may not be minimal. As a result, tree search was not restricted to the parsimony forest, but also trees derived from the parsimonious ones with a SPR approach. Heavy and light chains were kept separate for several reasons: (1) the mutation model differs between heavy and light chain^22^, (2) only one chain could be recovered for a substantial proportion of single cells (Supp. Fig.9A). SeQuoIA thus selects a tree pair with most consistent lineages between both chains, and penalises trees breaching current knowledge of BCR biology based on mutation load, productivity and CSR order (see Methods and Fig.2A for details). For each clonotype in an NP-KLH immunised mouse dataset^23^, we compared biology-based metrics between trees selected either with SeQuoIA or the original GcTree algorithm. GcTree could select trees with reverted mutations, while this criteria was systematically corrected with SeQuoIA (Fig.2B, Supp. Fig.9C,D). Aberrant parent to child relationships affected a larger number of cells in large clones (Fig.2B). Likewise, SeQuoIA’s ability to optimise tree pairs concordance was compared against tree pairs selected independently with GcTree. SeQuoIA could erase or decrease the number of aberrations in heavy/light chain lineages for most clonotypes (Fig.2C). One example of SeQuoIA correction is shown in Fig.2E. Among clonotypes with non-aberrant chain lineages, SeQuoIA further optimised lineage overlap in tree pairs (Fig.2D).

**Figure 2.**
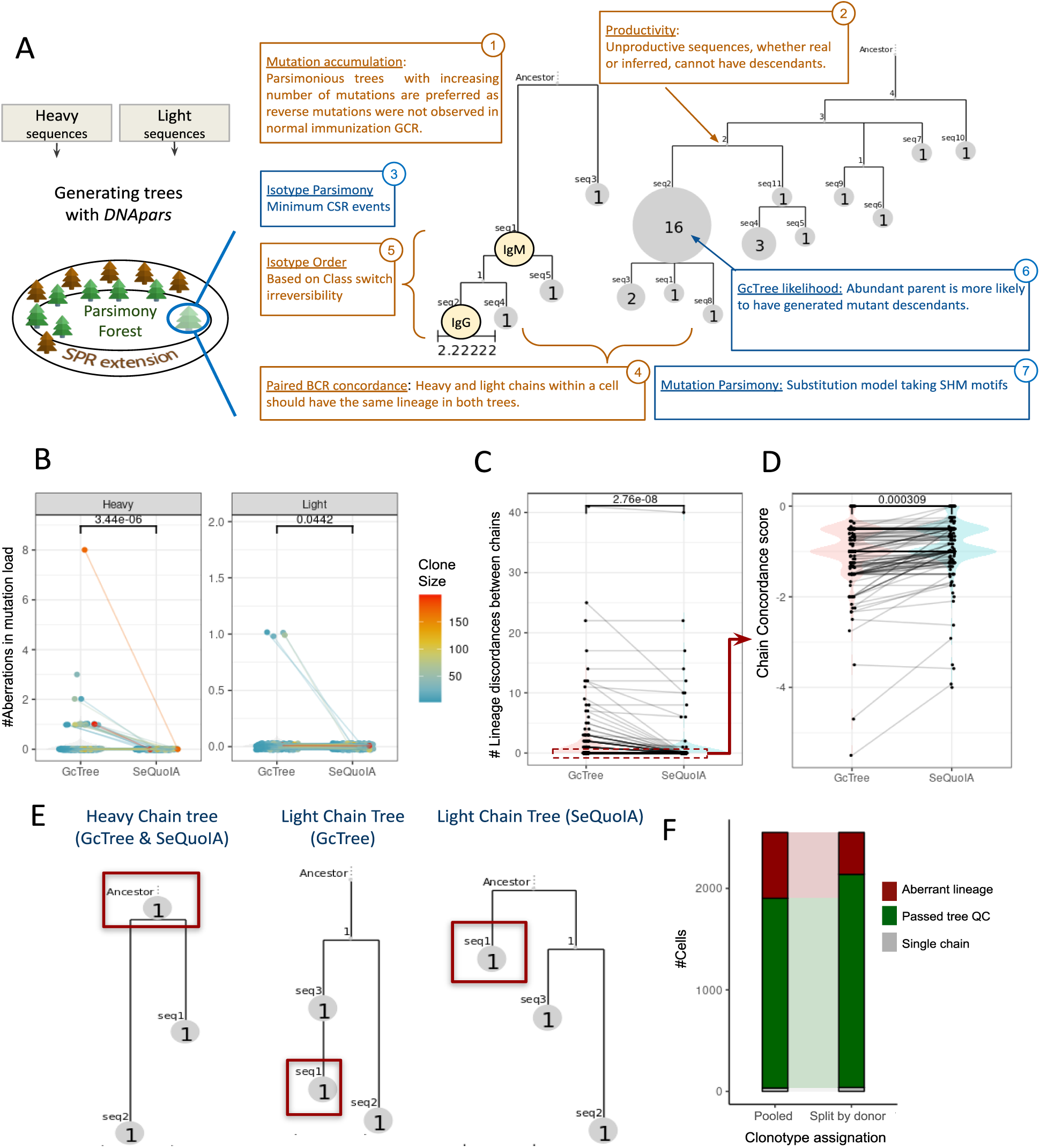
BCR phylogeny reconstruction in SeQuoIA. **A)** Schematic overview of phylogeny reconstruction. Parsimony method is applied for heavy and light chain phylogeny reconstruction independently, generating two tree exploration spaces. This space is extended thanks to the SPR approach. Several criteria (boxes) are then used to select one pair of tree from the heavy and light chain spaces. Purple boxes correspond to criteria already implemented in the GcTree package, while orange criteria are custom in SeQuoIA. Numbers in the boxes corners indicate the order in which the criteria are applied. **B)** Quantification of unexpected mutation loads (criterion n°1). Each dot corresponds to a single chain tree. **C)** Dot plot comparison of the number of incompatible tree pairs (criterion n°4) between SeQuoIA and GcTree. Each dot corresponds to a clonotype with paired heavy and light sequences. The p-value from a paired wilcoxon rank sum test is indicated. **D)** Dot plot comparison of chain concordance scores between SeQuoIA and GcTree for trees with no lineage discordance across chains. The p-value from a wilcoxon rank sum test is indicated. **E)** Example of lineage aberrance correction by SeQuoIA on a small clone. Red boxes correspond to a single cell, with incompatible positions in heavy and light chain trees in the GcTree solution. **F)** Bar graph comparison of tree reconstruction performance after two clonotype assignation strategies: considering all cells from the dataset in one analysis, or splitting cells by donor sample. Only clones with more than 5 distinct Igs were considered here to discard trivial trees. All analyses were performed on mouse NP-KLH1 dataset and GcTree was applied to heavy and light chains separately.

Despite exploring a larger space, SeQuoIA sometimes failed to find concordant tree pairs. We hypothesised these residual aberrations could stem from upstream steps in the pipeline and a propagation of errors in subsequent stages. Notably, the large size of some clones for which SeQuoIA did not converge to a solution raised the question as to whether they corresponded to a unique lineage descending from the same precursor. Clonotype accuracy was all the more questionable given that NP hapten immunisation is well known to elicit canonical variable segments (IGHV1-72/IGLλ, IGHV1–53). We thus checked whether such clonotypes corresponded to a single donor or were artefactual, by combining BCR data and donor data from the gene expression library. This analysis revealed that a substantial proportion of clonotypes were of mixed donor origin (Supp. Fig.10A). The share of each donor was balanced within artefactual clonotypes, thus ruling out technical noise from donor hashtag demultiplexing (Supp. Fig.10B). Largest clones (193 and 79 cells) did not correspond to single lineages, although small clones could also be inaccurate. Artefactual clones were not specific to the low diversity hapten response. Similar observations were made in polyclonal mouse response to influenza infection^3^ (Supp. Fig.10C-D). Conversely, clonotypes in the human tonsil dataset were not affected (data not shown). This divergence was further investigated with IGOr in Supp. Fig.4, which predicted higher recombination probabilities in murine clones. We sought to improve clonotype accuracy by incorporating donor origin in the assignment process. This option in the SeQuoIA pipeline enabled to decrease the number of aberrant lineages in phylogenetic trees (Fig.2F), suggesting that part of SeQuoIA’s failures to reconstruct concordant trees is due to the presence of unrelated cells within a clone when donor origin is not specified.

**Figure 3.**
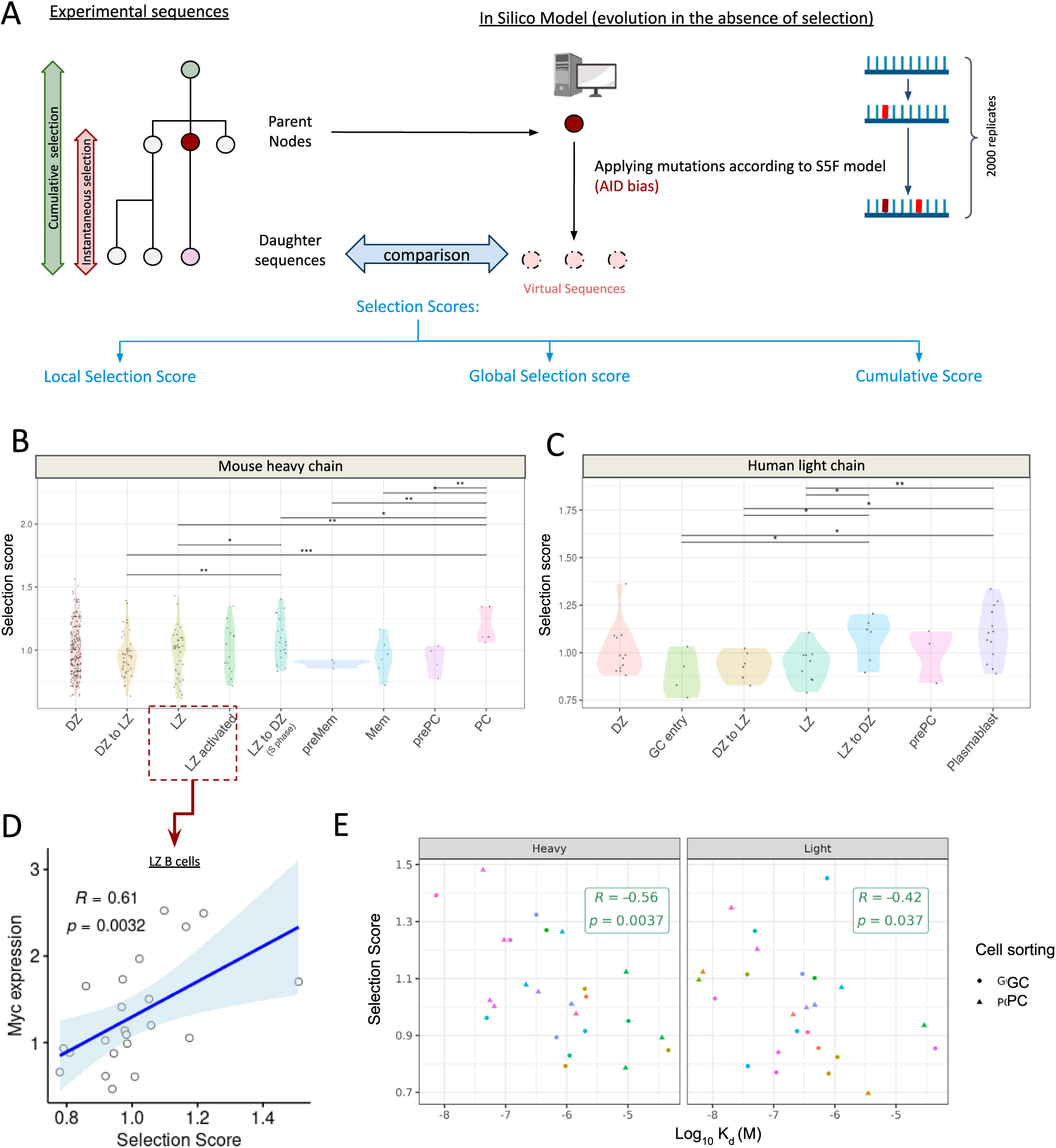
Inference of selection pressure in SeQuoIA. **A)** Schematic overview of the selection quantification part in SeQuoIA. Each circle corresponds to a single chain BCR sequence. Their relationships are represented by solid lines. **B)** Dot plot distribution of global selection scores in heavy chains across B cell subsets identified through scRNAseq cluster annotation in mouse NP-KLH1 dataset. Each dot is a cell. * p<0.05, ** p<0.01, *** p<0.001, non-adjusted one-sided wilcoxon tests. **C)** same as B in human tonsil dataset **D**) Spearman correlation of light chain BCR selection score and *Myc* expression for total LZ B cells in mouse NP-KLH1 dataset. Each dot corresponds to a cell. **E)** Spearman correlation between in vitro measured affinity to HA and heavy chain (left) or light chain (right) BCR selection score in mouse influenza dataset. Colours correspond to clonotypes.

**Figure 4.**
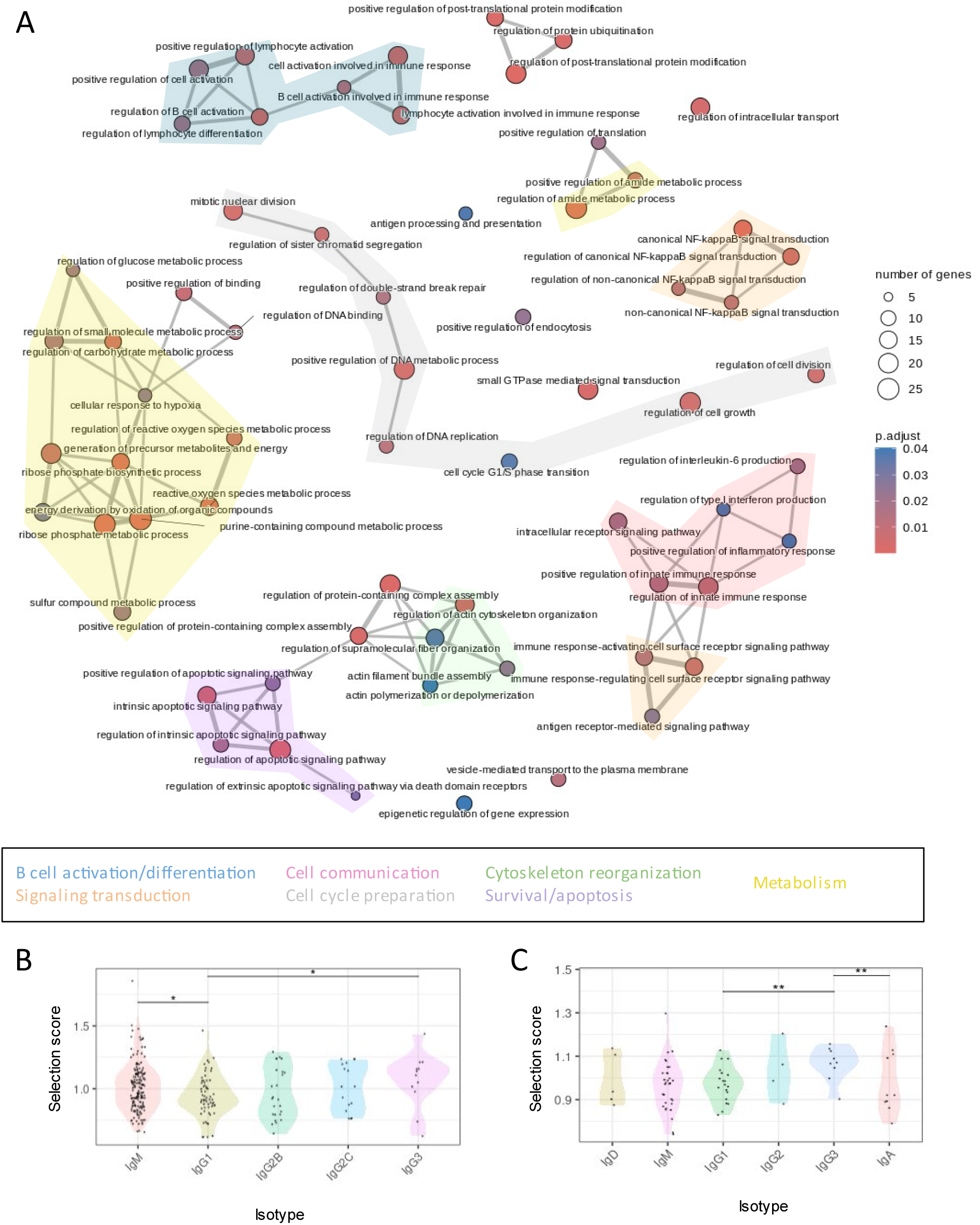
Application of SeQuoIA to find biological processes involved in affinity maturation in the LZ. **A)** Graph representation of Gene Ontology terms identified for genes whose expression was positively correlated to the light chain selection score among LZ B cells from mouse NP-KLH1 dataset. Nodes correspond to biological processes from the ClusterProfiler database and edges connect overlapping gene sets. B) Dot plot distribution of light chain selection scores for GC B cells grouped by isotype in mouse NP-KLH1 dataset **C)** Dot plot distribution of light chain selection scores for GC B cells grouped by isotype (entrants and GC B cells from secondary responses are excluded) in human tonsil. P values denote two sided wilcoxon tests, * p<0.05, ** p<0.01

### SeQuoIA selection inference and validation in public datasets

#### Inferred selection increases during LZ transit and correlates with Ig affinity

The substrate of selection is the accumulation of different somatic mutations in Ig encoding regions. SeQuoIA thus analyses mutation patterns to infer selection pressure based on their occurrence likelihood. Briefly, parental sequences (i.e. sampled internal nodes in the trees) seed a model of neutral BCR evolution. Mutations are applied according to the appropriate AID mutation model depending on species and Ig chain. Experimental and *in silico* generated sequences derived from the same parent are then compared and the divergence from non-selected virtual sequences is quantified with selection scores either at the codon or sequence level (Fig.3A). We chose to include Ig synonymous mutations in our analysis, although they are often presumed to be neutral, as they were shown to affect RNA stability and splicing and translation rates in other contexts^24,25^, and might affect BCR turnover in the DZ^12^.

SeQuoIA selection inference was validated against current knowledge of B selection during immune response. We identified cell subsets at different stages of B cell maturation in public integrative single-cell datasets^23,26^, through cluster-based reannotation (Supp. Fig.6-8), and applied SeQuoIA. Our pipeline measured an increase in selection scores between mutations accumulated prior to GC entry and within GCs, where affinity maturation takes place (Fig.3C, Supp. Fig.11A). In the extra-GC compartment, antigen-independent IgD^+^ Memory B cells which are thought to be activated through their TLRs^27,28^ displayed the lowest score of all human LN mutated B cell subsets (Supp. Fig.11A).

It was shown that GC selection primarily takes place in the LZ and that cognate T_FH_ signals may instruct B cell recycling to the DZ^29^. Selection imprint is thus expected to be stronger in cells transitioning from the LZ to the DZ. An increase in selection imprint was detected from the LZ entry to the LZ exit, in GCs from immunised mouse and human tonsil specimens or FNA biopsies (Fig.3B-C, Supp. Fig.11C). PCs were associated with higher selection scores, consistent with enhanced antigen binding affinity in immunisation response reported in previous publications^2,3^. Similar patterns in BCR selection imprint were observed for both heavy and light chains (Supp. Fig.12A, Supp. Fig.11D).

Another validation was performed at the gene level. Current monitoring of selected cells in the LZ rely on c-Myc expression^30^, which is controlled by synergistic BCR signalling and T_FH_ feedback^31^. Selection imprint was not only higher in *Myc*-positive cells as compared to *Myc*-negative LZ B cells (Supp. Fig.12D), but selection scores scaled with *Myc* expression levels in *Myc*-positive cells in mouse response to NP (Fig.3D) and human response to COVID vaccine (Supp. Fig.11D). The BCR signaling marker *NR4A1* (encoding for Nur77), also scaled with selection scores in human LNs. Therefore, SeQuoIA selection scores are relevant in the identification of B cells harbouring BCRs with enhanced signalling and/or Ag internalisation capacity.

Finally, we sought to determine whether SeQuoIA selection scores were a good indicator of Ig binding efficiency. BCRs generated in GCs induced by influenza infection were cloned *in vitro* and their equilibrium dissociation constants (Kd) for hemagglutinin (HA) binding were determined experimentally in a recent publication^3^. Heavy chain selection scores strongly correlated with measured affinity (1/Kd), regardless of the phenotype (Fig.3E). Another feature of SeQuoIA is the inference of selection pressure at the codon scale. Examples of this application can be found in Supp. Fig.2. One particular mutation is well described for its antigen binding enhancing effect after immunisation with 4-hydroxy-3-nitrophenyl acetyl (NP) hapten: W→L transition on the 33^rd^ codon (W33L) in the IGHV1-72 variable segment. This mutation was associated with a positive selection score. Because its maintenance was not associated with a significant negative selection score (Supp. Fig.2C), W33L monitoring at late GCR stages may not be informative on BCR selection.

#### Synonymous vs non synonymous mutations

The NS-only option of SeQuoIA was tested likewise and resulted in similar selection score distribution across cell subsets (Supp. Fig.12B). However, this analysis led to a loss in statistical power due to fewer observations. Differences between DZ-to-LZ and LZ selection scores distribution were also less marked, suggesting a role for synonymous mutations in DZ to LZ transition. We next examined synonymous to NS ratios and respective scores across GC phenotypes. This analysis uncovered an enrichment in synonymous mutations after DZ exit (Supp. Fig.12C), while most mutations in the neutral evolution model are NS (Supp. Fig.13A). Synonymous mutations had intermediate to high selection scores, while a cluster of NS mutations displayed low selection scores. Altogether, this suggests that most mutations are deleterious even though Ig sequence is still productive and are purged upon LZ entry. Synonymous mutations, albeit unlikely, could be more prone to pass this putative DZ-to-LZ checkpoint, as they would not lead to decreased Ig fitness. Conversely, NS mutations were enriched in the LZ-to-DZ compartment, suggesting a more stringent selection step.

#### Comparison with BaseLine

Previous attempts of selection quantification based on somatic mutation patterns have been published^7^ and integrated within the *Immcantation* suite as the BaseLine tool. BaseLine compares replacement to silent mutation ratio from observed sequences and theoretical values from a bayesian estimator based on a binomial likelihood function and β prior. With this assumed distribution, the greater the mutation load, the smaller the confidence interval of the selection strength estimate. SeQuoIA, on the other hand, does not necessarily assume that silent mutations are neutral, and minimises the number of mutations for observed and theoretical sequence comparison. The phylogenetic reconstruction upstream of selection quantification allows to narrow down the modelling time scale to a few mutations (Supp. Fig.3F), thus limiting uncertainty on re-mutation and mutation hotspots/coldspots reshuffles. Our quantification method does not rely on replacement to silent ratio, which allows us to quantify selection at the codon level or to individual sequences without aggregation^7^. Furthermore, replacement probability is equal or close to 1 in most of modelled mutations (Supp. Fig.13A), which could lead to decreased ability to detect enrichment in NS mutation as compared to the model. Accordingly, when we applied *BaseLine* on the same dataset as Fig.3B, no significant difference was found between B cells entering or exiting the LZ (Supp. Fig.13B).

### SeQuoIA usage to investigate selection mechanisms in the GC LZ

Having validated our selection quantification approach, we next attempted to identify new LZ B cell selection mechanisms with our tool. To identify transcriptional programs elicited in cells expressing selected BCRs, we set up a gene screening approach in LZ B cells, where normalised gene expression was correlated to Ig selection score at the single-cell level. Candidate genes (spearman correlation coefficient > 0.2, p < 0.05) were submitted to Gene Ontology (GO) analysis to interrogate their functional relevance. Biological processes (BP) linked to Ig selection included expected cellular responses to B cell stimulation, such as *NF*_κ_B signal transduction which was shown to be triggered by CD40 stimulation in a subset of LZ B cell^31^, Ag receptor signalling, and Ag presentation (Fig.4A). At the gene level, *Irf4* expression, which was shown to be induced in response to a combination of high affinity antigen and T_FH_-derived cues in GC B cells^32^, displayed a strong correlation to the selection score (Supp. Fig.14). BCR signalling may also synergize or sensitise to interferons in the LZ, as *Stat2* levels were correlated to selection scores (Supp. Fig.14). Overall, SeQuoIA was able to identify activated LZ B cells involved in recent Ag binding events and interactions with T cells. Another relevant BP was regulation of cell cycle. Altering T_FH_ feedback to LZ GC B cells in *in vivo* mouse response altered GC size and cell division in the DZ, suggesting that BCR-mediated selection in the LZ regulates the proliferative capacity of GC B cells^33^. In line with these observations, the expression of regulators of cell cycle scaled with SeQuoIA selection scores (Fig.4A, Supp. Fig.14), supporting BCR-based selection in the LZ as a prerequisite for cell cycle progression.

Actin cytoskeleton organisation is also one of the BP worth mentioning; *in vitro* assays revealed that LZ B cells were able to discriminate high and low affinity antigens independently of cell to cell competition thanks to pulling forces exerted by their contractile cytoskeleton^34^. This discrimination resulted in affinity-dependent Ag internalisation^32^. Actin dynamics can also modulate BCR signalling capacity through BCR microcluster formation^35^. SeQuoIA disclosed BCR-related upregulation of actin cytoskeleton remodelling or vesicle-forming proteins (Supp. Fig.14), which supports the existence of a positive feedback at the transcriptional level where higher affinity BCR would benefit from stronger signalling transduction cascade in subsequent interactions and increased B-T synapse formation capacity.

Finally, selected BCR features seem to confer a competitive advantage to LZ B cells via enhancing resistance to apoptosis. BCR-driven selection was predicted to occur partly through the inhibition of pro-apoptotic factors from the BCL2 family (Supp. Fig.14). Interestingly, protection from the extrinsic pathway was also predicted to be fostered by selected mutations (Fig.4A), suggesting that LZ B cells with unfit BCRs could be actively eliminated instead of dying by neglect. Mechanistically, this protection would be conferred by the recruitment of caspase 8 inhibitors (e.g. FAIM, encoded by *Faim1*) in the Fas associated death domain FADD (Supp. Fig.14).

### BCR isotypes exhibit different selection strengths

In addition to Ig variable part diversifiable through recombination and SHM, BCRs have different cytoplasmic domains depending on their isotypes. These structural features could lead to different signalling capacities, mechanical force sensing or co-receptor interaction, thus conferring additional competitive advantage on top of affinity for the Ag. Notably, IgM BCRs were shown to have a higher activation threshold as compared to IgG1 BCRs in a biophysical *in vitro* assay^36^, and were outcompeted in *in vivo* GCRs^11^. In keeping with these observations, SeQuoIA inferred decreased selection pressure in IgG1 GC B cells variable domain, which could be possible through a “native” competitive advantage (Fig.4B). Another interesting observation was the increased selection imprint in IgG3 BCRs from both immunised mouse LN and human tonsil (Fig.4B-C). IgG3 may thus have attenuated signalling capacity, which would put an additional constraint on this population.

### Selection impact after LZ exit

#### DZ B cells

Unlike c-Myc expression reporters, which can only detect selected cells among LZ GC B cells, Ig mutation monitoring with SeQuoIA allows insight into selection strength impact on B cell phenotype beyond LZ exit, either as recycling DZ B cell or effector B cell. To investigate whether and how LZ selection determines GC B cell fate once they enter the DZ, we set up a new correlation analysis relying on phylogenetic tree knowledge. Briefly, genes expressed in the DZ were screened for correlation between gene expression levels and parental LZ selection score, if sampled (Fig.5A). LZ parent and DZ progeny pairs were retained if their BCR differed by less than 2 mutations, to restrict the analysis to direct parent-progeny relationships. A GO analysis was run next as described previously (Fig.5B). Among the BPs linked to LZ selection, SeQuoIA found DNA biosynthesis and cell cycle regulation, in accordance with previous study concluding in accelerated S phase progression for DZ B cells after enhancing T_FH_ feedback in the LZ^37^. Additionally, response to DNA damage and DNA repair emerged from this analysis, which could be consistent with increased SHM mutation in selected cells. On the other hand, there was no correlation between LZ selection score and AID or error prone Polη expression in DZ descendance (data not shown). CXCR4 expression scaled with LZ parent selection score, suggesting that fit GC B cells could benefit from a DZ stromal cell niche, or stay longer in the DZ, as shown in photoactivation assays of DZ B cells^38^. In turn, extended DZ residence time could lead to an increased number of divisions and somatic mutations. GO analysis also revealed the possible involvement of a transcriptional shift in metabolic pathways (Fig.5B), including the upregulation of fatty acid transport (*FABP5*), and mitochondrial ATP synthase and oxidoreductases (*ATP5F1A/SCD1*). The upregulation of genes involved in oxidative phosphorylation in the DZ seems to emanate from BCR-related expression of regulators of metabolism (amide and carbohydrate) and the generation of precursors in the LZ (Fig.4A). The regulation by LZ cues of subsequent DZ metabolism supports previous observation of increased OXPHOS related gene expression in B cells carrying affinity enhancing mutation W33L in the response to NP antigen^39^. Finally, inositol phosphatase *INPP5B* was strongly correlated to LZ selection score (Fig.5C). This phosphatase is involved in inositol triphosphate inactivation, and could thus terminate calcium mobilisation downstream of BCR signalling. This branch of BCR signalling was shown to be involved in a negative feedback, with notably the upregulation of inhibitory coreceptor *CD72*^40^.

**Figure 5.**
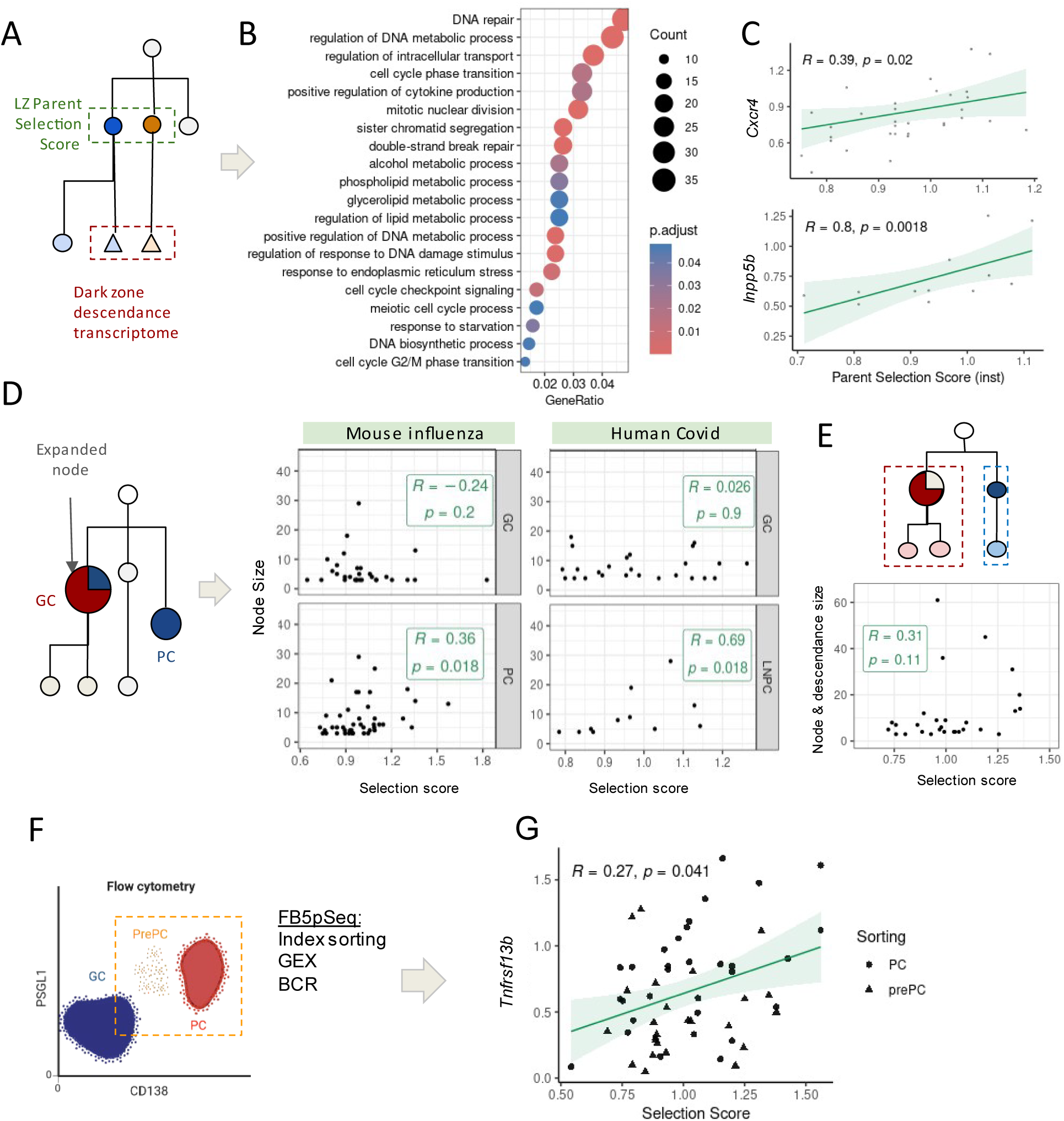
Application of SeQuoIA to investigate the impact of selection after LZ exit **A)** Schematic overview of the analysis. DZ B cell gene expression was correlated to its LZ parent (if sampled) selection score. This analysis was performed on mouse NP-KLH1 dataset. **B)** Significant Gene Ontology terms from analyzing genes in DZ progeny with significant correlation (ρ > 0.2, p.value < 0.05) with LZ light chain selection score **C)** Scatter plot and Spearman correlation of the expression of selected genes in DZ B cells and light chain BCR selection score in their LZ parent. **D)** Schematic overview and analysis of Spearman correlation between node size and its associated heavy chain selection score, with nodes split by GC and PC cell types. Only expanded nodes (i.e. > 3 sequences) were considered. This analysis was performed in in mouse influenza and human covid datasets. **E)** Schematic overview and analysis of Spearman correlation between node plus progeny size and the node’s light chain selection score, focusing on GC B cells in mouse influenza dataset. **F)** Schematic of mouse PC and prePC enrichment experiment from mouse NP-KLH2 dataset **G)** Spearman correlation of prePC and PC light chain selection scores with *Tnfrsf13b* gene expression.

#### Proliferation rates

As discussed in the previous section, reduced cell death in the LZ seems to contribute greatly to the competitive advantage of selected cells and BCR-based selection may be a prerequisite to cell cycle entry. Whether increased cell division from fit cells also contributes to their maintenance over the GCR has already been hypothesised^41^ but not demonstrated in untouched polyclonal responses to complex antigens to our knowledge. On top of GC “bursts”, recent GC-derived PCs were shown to proliferate upon GC exit^23,3^. The decoupling between DZ and PC proliferation raises the question as to whether one of them is driven by BCR-mediated selection. We thus reanalyzed Sprumont et al dataset and compared PC and GC tree node sizes correlations to their selection scores. Of note, nodes were split if a same node was associated with multiple phenotypes. PC selection scores, unlike those of GC B cells, were mildly correlated to the node size (Fig.5D), suggesting that BCR features may determine, at least partly, the extent of post-GC proliferation. This pattern was also observed in human response to SARS-Cov2 vaccination (Supp. Fig.15A). However, it could be argued that somatic mutations are a confounding factor as they often result in selection score decrease and co-occur with DZ proliferation. Hence, another correlation was drawn between SeQuoIA selection scores and GC node size and direct descendance, which was not significant (Fig.5E). Altogether, these results suggest some stochasticity in GC proliferation, whereas PC expansion (or survival) would be somehow linked to Ig features.

#### PC differentiation

PC differentiation is another fate after transiting through the LZ, which has been a focus of numerous publications^2,3,42^ since PCs are at the frontline in the fight against pathogens. This compartment was reported to be enriched in Ag binding cells as compared to total GC compartment in mice immunised with NP^43^ or HEL^2^ in past and recent literature. On the other hand, low affinities were measured in BCRs harboured by GC-derived MBCs^44^. SeQuoIA selection scores were measured in GC and output cells in murine (Fig.3B, Supp. Fig.12A) and human tonsillar or LN aspirate datasets (Fig.3C, Supp. Fig.15C). In all these datasets, in light and heavy chains (if available), selection imprint was lower in MBCs as compared to LZ B cells, whereas PCs displayed higher selection scores, consistent with enhanced affinity reported previously^2,43^.

Nevertheless, recent publications suggest that PC differentiation could be more permissive to low-affinity cells than anticipated. In polyclonal GC responses to plasmodium and influenza infections, PC precursors^45^ and PCs^3^ displayed similar affinities as GC B cells. This recent literature may seem at odds with stringent selection inferred with SeQuoIA in four independent datasets. However, this selection imprint could be affinity-independent, including autoreactive prePC/PC elimination^46^. Since the nature of selection cannot be deconvoluted with SeQuoIA, we tried different strategies to further investigate PC selection in complex human and above-mentioned murine responses datasets. In agreement with Sprumont et al.^3^ affinity measurements, unique BCRs from PC compartments had similar selection scores as GC B cells (Supp. Fig.15A). However, global PC population displayed an increased selection imprint thanks to expanded selected cells (Supp. Fig.15B), suggesting that selection operates in the PC compartment through differential proliferation, possibly in an affinity regulated manner^23^.

Conversely, a shift in selection scores between GC B cells and PCs was observed on unique sequences from the human covid dataset (Supp. Fig.11D). We had similar findings in human tonsil dataset (Fig.3C), comprising small clones without expanded nodes. Selection into or within PC compartment may thus not be restricted to proliferative effect. Divergence between results obtained on human datasets and polyclonal murine model could be explained by (1) exclusion of non-binding cells through cell sorting (2) FTY720 treatment preventing PCs from egressing LNs, possibly confounding past and present PC generation. Within this line, selection imprint increased in both GC and PCs within a few days (Supp. Fig.15C), suggesting that selection imprint distribution in influenza dataset might have been partly brought down by earlier-generated PCs.

We thus investigated in untouched GC responses whether PC differentiation could be subject to selective pressure. Interestingly, cells having an intermediate phenotype between GC and PCs, henceforth referred to as “prePC”, displayed lower selection scores than PCs and putative late DZ PC precursors (Fig.3B-C, Supp. Fig.11A, Supp. Fig.11C). Most often, they had equivalent selection scores to LZ B cells. Altogether, increasing selection score distribution over PC differentiation trajectory suggests that a selection step could occur during and/or after the prePC to PC transition. Unique PC Ig sequences displayed higher selection scores as compared to prePC (as well as LZ B cells) regardless of post GC expansion (Fig.3C, Supp. Fig11C), suggesting that prePC to PC transition is not only subject to positive selection through proliferative advantage. Negative selection could clear unfit prePC or positive selection could licence fit prePC into becoming fully fledged PCs.

We therefore analyzed prePC and PC gene expression correlations with SeQuoIA selection scores to identify putative competitive advantages conferred by Igs during PC differentiation. This analysis was carried out in a published dataset^23^ where PCs and prePCs were enriched prior to scRNAseq (Fig.5D). We found significant correlation between Ig selection score and *Tnfrsf13b* (Fig.5E), suggesting that late stages of PC differentiation are linked to BCR-driven selection. APRIL signaling through the TACI receptor, encoded by *Tnfrsf13b*, has been shown to support PC differentiation upon GC exit at the GC/medullary interface (GMI)^47^. We also identified other relevant genes such as *Cd37* (Supp. Fig.15D), a marker of long-lived plasma cells (LLPCs)^48^. Anti-apoptotic factor *Caap1* also turned out to be correlated to Ig selection (Supp. Fig.15D), indicating that a competitive advantage in terms of survival may be conferred to prePCs and PCs during or after differentiation. Finally, cell growth regulator *Mtor* scaled with SeQuoIA selection scores (Supp. Fig.15D), supporting results from the previous section where PC numbers correlated with their Igs selection imprint. All in all, Ig features seem to determine PC fate during and after their differentiation, in terms of survival, proliferation and possibly BM homing.

### Selection pressure evolution over time

GCs are highly dynamic structures with variable lifetime. Despite the fact that individual GCs are asynchronous^49^, several mechanisms may modify the selection pressure over the course of GCRs, including the emergence of regulatory GC T cell population^50^, a switch from MBC-to PC-dominant exportation regime^42^, or increasing serum antibody affinity resulting in tighter antigen retention at the FDC surface^51^. To assess whether selection pressure evolved over time in GCRs, we took advantage of a published dataset^23^ consisting in mouse LN sampling at different timepoints after immunisation (Fig.6A). SeQuoIA detected increased selection imprint in GC B cell heavy chains over time post-immunisation (Fig.6B). Temporal changes in selection score were also correlated to the mutation load in GC B cell light chains, suggesting that selection strength relies on time spent in GCs (Fig.6C). Finally, SeQuoIA was applied to a clinical cohort of SARS-CoV2 vaccination to assess time required for immune protection (Fig.6D). Selection scores were still low at day 60 after primary vaccine injection, but increased between day 60 and day 201 post vaccination, suggesting that effective and long-lasting affinity maturation was elicited by the vaccine after a phase of relative latency in selection (Fig.6E).

**Figure 6.**
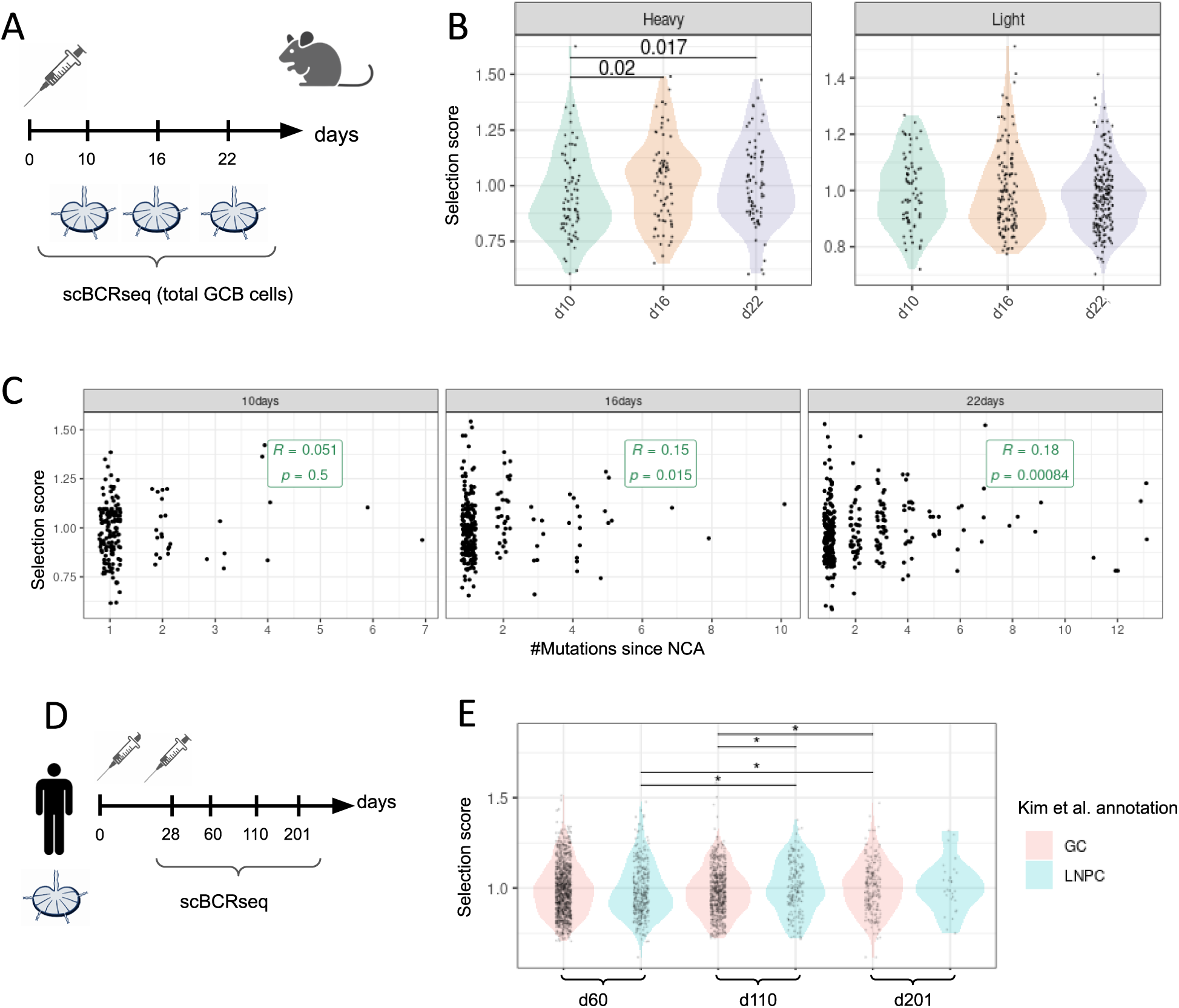
Evolution of selection pressure in germinal centres over time. **A)** Schematic overview of immunization and sampling from mouse NP-KLH3 dataset. Cells from draining lymph nodes of two mice were analyzed at each timepoint. **B)** Dot plot analysis of selection scores in IgH (left) and IgL (right) at different time points after immunization. Each dot represents a sequence 1 mutation away from its parent. p-values denoting Wilcoxon rank sum test results are indicated. No multiple testing correction. **C)** Scatter plot of selection score by number of mutations from clonotype NCA (considered as BCR phylogenetic tree pseudotime) in the light chain at given time points after immunization. The selection score is significantly correlated (Spearman correlation) to distance from tree root at intermediate and late timepoints. **D)** Schematic overview of vaccination and sample collection time points in human Covid dataset**. E)** Dot plot distribution of selection scores in GC B cells (pink) and PC (green) at the indicated time points after vaccination. *p<0.05, wilcoxon rank sum test, non-adjusted.

## Discussion

We herein introduced a start-to-finish pipeline for single-cell Ig repertoire analysis in clonally expanded B cells, with improvements of current clonotype assignment and Ig tree reconstruction methods. Most importantly, we developed a modelling approach for BCR-driven selection inference.

To date, BCR-based selection in GCs has been overwhelmingly studied through a few mouse immunisation models with small antigens (NP, HEL), which are characterised by stereotypical BCRs and short-lived GCs, and the monitoring of a handful of recurrent mutations. Although those models have proved valuable to explore basic concepts of GC function in controlled settings, our analyses with SeQuoIA highlighted some of the limitations of such models: low clonotype assignment accuracy, low pre-GC selection, irrelevance of W33L mutation monitoring at late stages. SeQuoIA, on the other hand, opens up new avenues for studying competitive advantage conferred by selected mutations, including in complex responses and human samples. Taking into account the light chain not only increased phylogeny reconstruction accuracy, as assessed by others in trees of concatenated heavy and light sequences^52^, but was also useful for detecting selection patterns. The increased availability and low mutation load since the last parent proved to be valuable, especially when screening gene expression for correlations to the selection scores. These selection scores were validated against affinity measurements and against the expression of known activation markers (NFκB, MYC, IRF4).

Having validated our method, new mechanisms for BCR-driven selection were put forward. We propose BCR signalling capacity as being an integral part of the LZ selection process, that could be a first checkpoint allowing the acquisition of LZ phenotype (Supp. Fig.16). A shift in selection score distribution was indeed detected in productive cells from DZ-to-LZ to LZ phenotype. NS mutations seemed to be purged in late DZ stages, before DZ-to-LZ transition. Altogether, these observations suggest that failure to reach the LZ or acquire LZ phenotype is not solely due to an absence of BCR or tonic signalling. One mechanism for this first checkpoint can be found in the Luo et al. study which demonstrated that BCR stimulation is sufficient for Foxo1 translocation from nucleus to cytoplasm, thus allowing the DZ to LZ switch^31^.

Once in the LZ, the screening approach identified resistance to apoptosis (intrinsic and extrinsic) as a transcriptional pattern associated with BCR-driven selection. Selection in the LZ is thus likely not to be reduced to competition for limiting factors and death by neglect of outcompeted B cells, as stated in previous work^1^, especially as FAS-mediated cell death was shown to be involved in autoreactive and bystander GC B cell elimination^46^. Changes in FADD composition in connection with BCR properties provides one mechanism explaining the protective effect conferred by BCR signalling regarding FAS-mediated cell death which has been reported in early experiments^53^, but remained unelucidated so far. Besides the competitive advantage in terms of survival, the expression levels of membrane proteins involved in B/T synapse were correlated to SeQuoIA selection scores. BCR signalling may thus influence T_FH_ help beyond antigen internalization, through transcriptional regulation of surface receptors. SeQuoIA also identified regulation of metabolic programs which then translated into differential expression of enzymes/transporters connected with OXPHOS in the DZ progeny. This metabolic shift may support DZ B cells survival and proliferation and be directly or indirectly regulated by BCR binding properties.

SeQuoIA did not find a correlation between BCR fitness and GC B cells abundance, but genes involved in the regulation of cell cycle seemed to licence fit cells to undergo cell divisions. The link between BCR properties and proliferation may be blurred by concomitant accumulation of somatic mutations (most of which are deleterious), or a stochastic component. Stochasticity in proliferation rates may lie in cognate T_FH_ encounter, which may have not been captured in current mouse studies resorting to anti-DEC-205-mediated antigen delivery and antigen-specific transgenic T cell transfer^33^. If differential proliferation did not seem to be elicited by BCR selective value, cell death in the LZ appears as a major driver of BCR-driven selection.

Finally, selection imprint was found to evolve along the GCR and immunisation/vaccine time. Several putative mechanisms for increased selection pressure over time were also provided in this study, including the aforementioned calcium-dependent BCR negative feedback in unfit cells and enhanced BCR-signalling and Ag presentation capacity through cytoskeleton remodelling in fit cells. BCR-dependent phosphorylation of actin nucleators was already demonstrated^35,54^, thus altering cytoskeleton through post translational regulation, but SeQuoIA application to *in vivo* datasets revealed the possible involvement of a long term bidirectional feedback loop with modulation of cytoskeleton composition through BCR-dependent transcriptional regulation.

The last application of SeQuoIA provided insight into the selection events leading to PC generation. We propose a model that could partly reconcile diverging literature on PC selection^3,45^. Our results are compatible with Sutton et al study^45^, where no enrichment in affinity enhancing mutations was found in prePCs (at least when they still have a LZ phenotype). We identified a putative new BCR selection checkpoint during or after prePC to PC transition upon GC exit, explaining increased affinity (or autoimmune cell clearance) in the PC compartment (Supp. Fig.16). PrePC-to-PC selection checkpoint could control maturation, local PC proliferation rates, and PC fate once they exit GCs and LNs, such as their longevity. The latter falls in line with a publication^55^ where LLPC numbers were not affected by the different amounts of newcomers in the BM niche, suggesting that the turnover is determined, at least partly, by intrinsic properties. We identified anti-apoptotic Caap1 as a possible determinant factor of LLPC lifetime, which expression would be settled upon GC exit according to the Ig properties. Whether the Ig related competitive advantage conferred to some prePCs and LLPCs is determined by upstream LZ cues or by residual surface BCR expression^56–58^ warrants further investigation (see Supp. Fig.16).

The extensive usage of SeQuoIA in this study to investigate GC selection mechanisms is a foretaste of its potential applications. For instance, whether the different selection imprints in GC B cells harbouring different isotypes is linked to intrinsic BCR properties or is dependent on the cytokinic context has not been explored. We hope that this pipeline could have applications to study autoimmune or lymphoma BCR-related pathogenesis, to assess B-cell responses in tumour tertiary lymphoid structures. Clinical applications could include patient stratification in the aforementioned pathologies, monoclonal antibody design and vaccine response assessment.

### Limitations of the pipeline and the study

We would like to stress that the pipeline has no application in datasets restricted to non-clonally expanded B cells (*e.g.* circulating B cells) since clonal phylogenies are required for inferring selection pressure. Despite the robustness of SeQuoIA selection scores, some Ig repertoire analysis issues have not been resolved and the application of this pipeline may be limited by the computation time and computational resources usage. Applications in murine datasets may be less reliable than in human datasets, since clonotypes (as classically defined) may not be fully accurate. Furthermore, the murine mutation model that we built is likely to lead to an underestimation of selection imprint as it was derived from spontaneous GCs in PPs, where GC B cells should nonetheless still be subject to self-tolerance. As our donor-based clonotype assignment did not fully resolve the quality of phylogeny reconstruction, residual intra-donor chimeric clones or uncorrected sequencing errors could explain the absence of solution following tree space exploration. Finally, mutation gaps in trees and low overlap between GEX data and post-filtering Ig data lead to decreased statistical power. Indeed, our results proved robust as they were convergent in four independent human and murine setups but the sensitivity of our tests was limited by the low number of informative cells (cells with complete gene expression and BCR sequencing data, no mutation gap). Within this line, one major limitation of this study is that the most abundant clonotype of each dataset had to be discarded from the analysis due to tree reconstruction errors. We are nevertheless confident that improvements in sequencing technologies and BCRseq preprocessing would partly increase the overlap between informative BCR sequencing and gene expression data. Experimental strategies could optimise the amount of analysable data, especially in human datasets, such as small tissue wedge dissection, which allowed recovery of larger clonal portions as compared to whole tissue processing in previous publication^59^. Finally, we were not always able to perform gene screening on both chains. We hypothesise that the ability of SeQuoIA to correlate selection scores and gene expression levels depends on the chain transcript availability, antigen type, and evolution speed as there could be some latency for last mutation to translate into gene expression modulation.

## Code & data availability

The SeQuoIA source code and dockers are available on Zenodo. All information and a tutorial can be found on GitHub (https://github.com/MilpiedLab/SeQuoIA). All datasets used in this paper are summarised in Supp. Table 3.

## Acknowledgments

This work would not have come to fruition without the help of Romain Fenouil for docker environments troubleshooting, including the containerization and adaptation of GcTree to the pipeline. Sabrina Baaklini’s help on the onset of the project was highly valuable with her pioneering work in heavy and light chain clonotype assignment. We thank Thierry Mora for his support and the profitable discussions that led to the current sequence selection score. We acknowledge the Computational Biology, Biostatistics and Modeling (CB2M) hub at CIML for setting the computational working environment and the discussions on the pipeline availability and reproducibility. We are also grateful to Chuang Dong for the datasets handover. We thank past and current members of the “Integrative B cell immunology” lab at CIML for fruitful discussions on GC selection mechanisms. Finally, we would like to thank Will Dumm, Erik Matsen, Adam Ye, Frederick Alt, Ali Ellebedy, Adrien Sprumont and Oliver Bannard for their availability and for kindly providing code and data information. This work was supported by a PhD scholarship to EH from the French Ministry of Research and Higher Education. This work was supported by the Fondation pour la Recherche Médicale, grant number FDT202404018734, to EH. This work was supported by a grant from Agence Nationale pour la Recherche (projet “GCselection”, ANR-23-CE15-0025-01) to PM, and by institutional grants from INSERM, CNRS, and Aix-Marseille University to the CIML.

## Supplementary Tables and Figures

**Supplementary Table 1.**
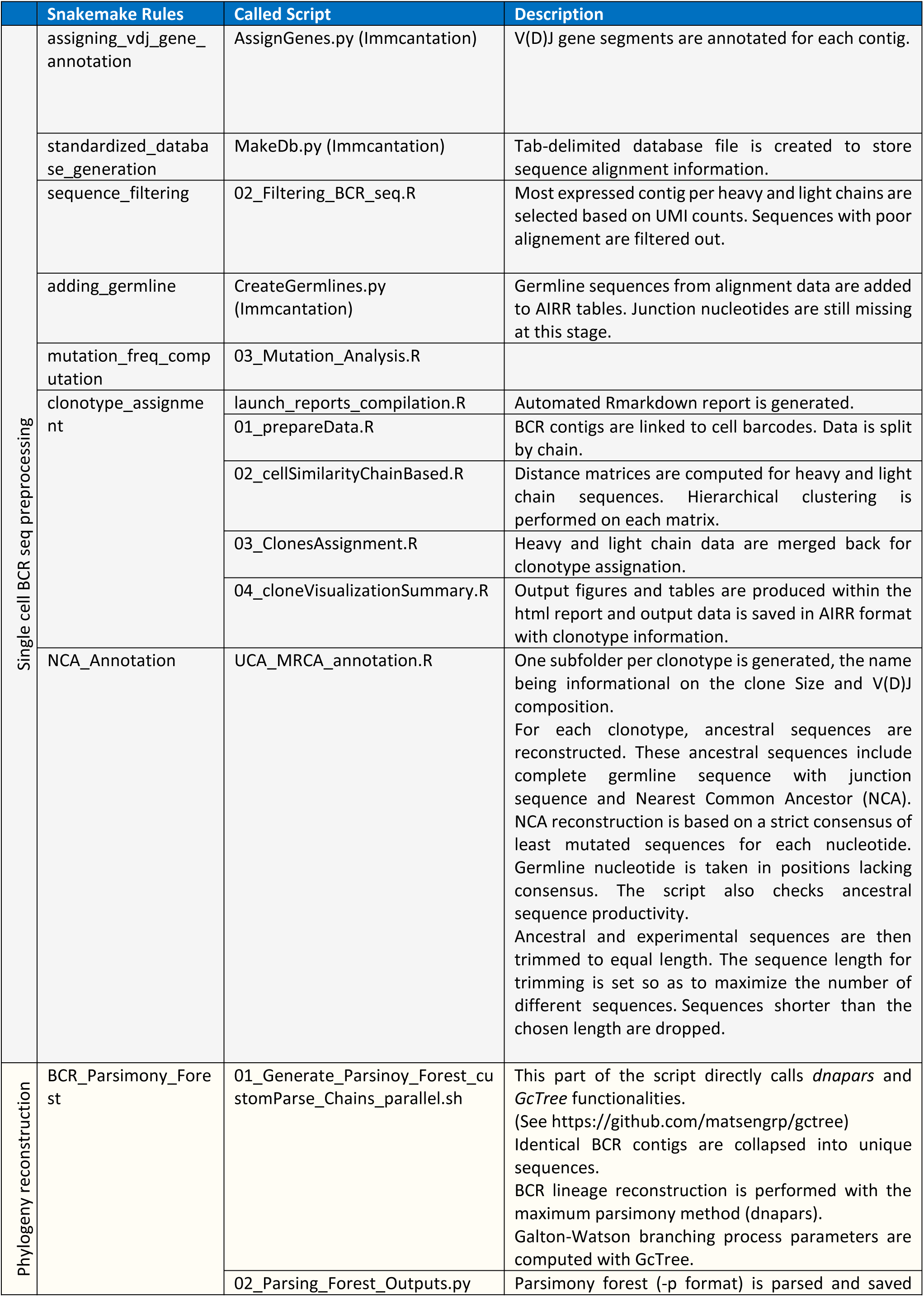
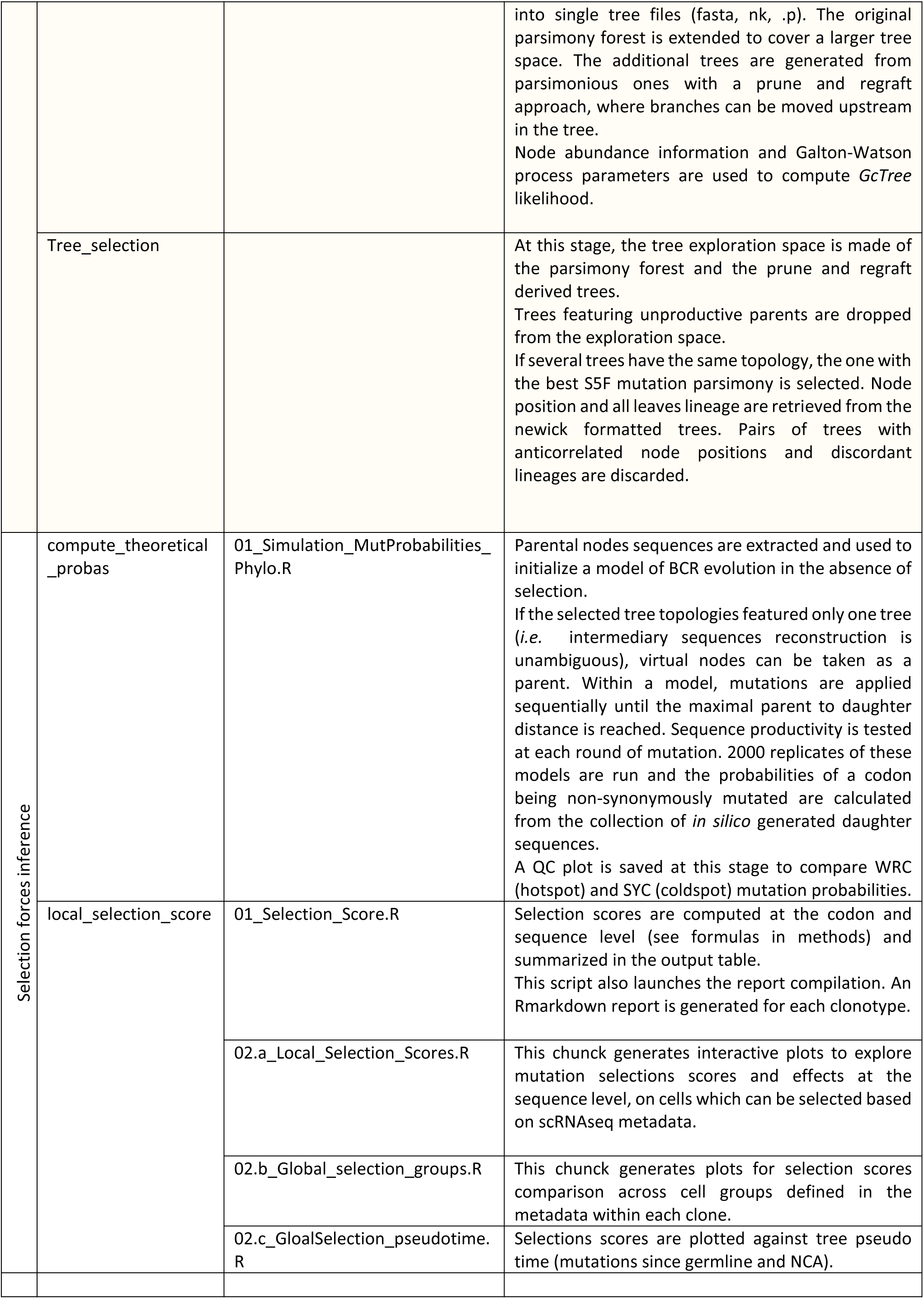
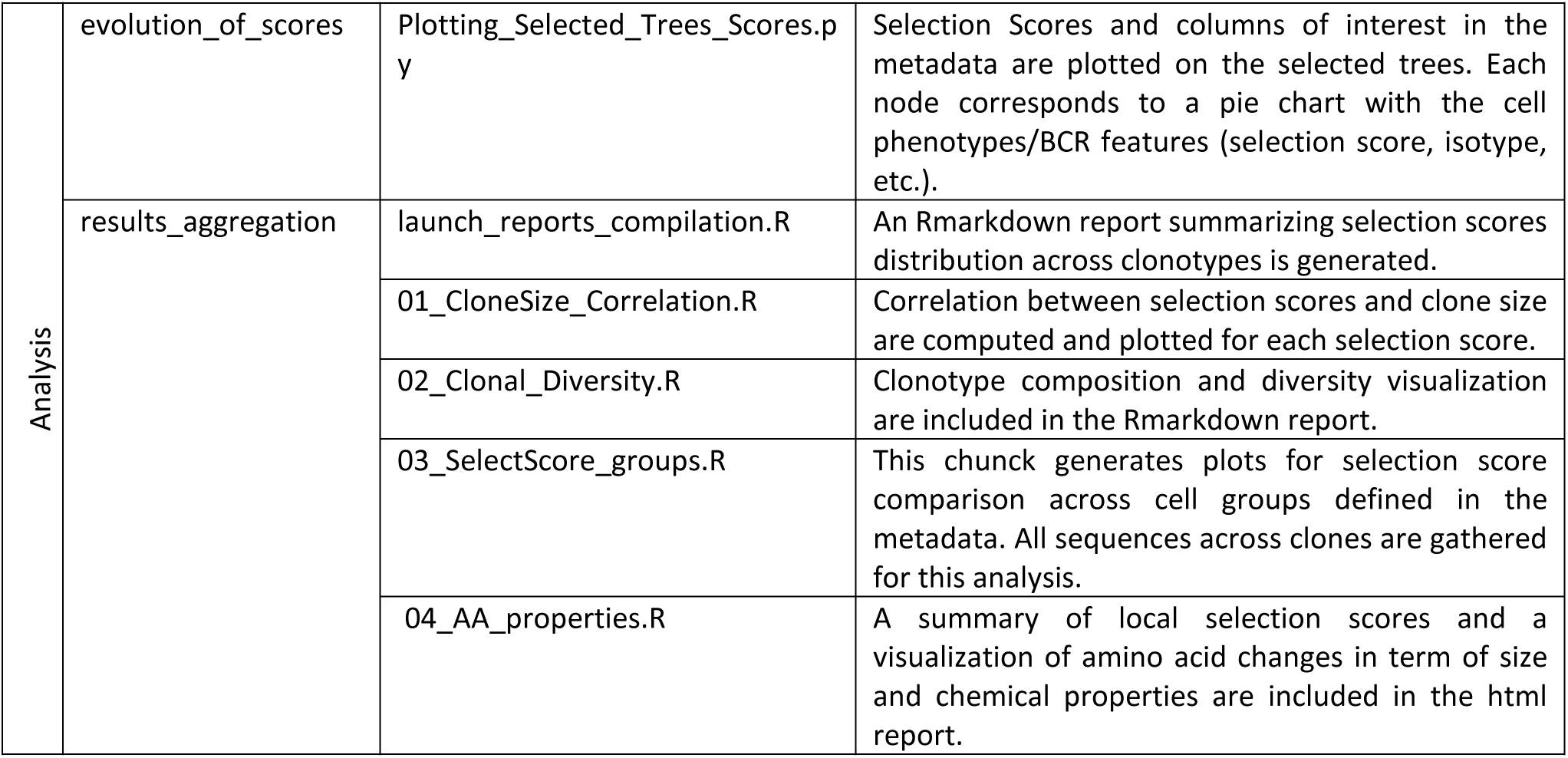
Overview of scripts in the SeQuoIA pipeline (related to Fig.1).

**Supplementary Table 2.** Overview of data inputs into the *SeQuoIA* pipeline (related to Fig.1).

| Name | Localization | User involvement | Format | Usage |
| --- | --- | --- | --- | --- |
| *.fasta | Rawdata > VDJ | Mandatory | > BCR_contig_id sequence | experimental sequences |
| *annotation.csv | Rawdata > VDJ | Mandatory | cell ranger vdj-like format. cell_id contig_id columns are mandatory. | Linking chain sequences to single cell id |
| *metadata*.extension | Rawdata > scRNAseq | Optional but highly recommended | txt/csv accepted must contain "barcode" or "cell_id" column. Otherwise, rownames will be taken as cell ids. | Comparison of selection scores between groups of cells (annotated during the analysis of scRNAseq data). |
| config_BCRseq.yaml | Workflow | Pre-filled. Species variable is mandatory. | fields are pre-filled but are editable:<br>-computational resources<br>-species (mouse or human)<br>-metadata column of interests (optional) | computational resources for parallelization, species for alignment, relevant metadata |
| mouse and human reference genomes | Reference > Genomes Alignments | Built-in but editable | fasta | germline sequences for BCRseq alignment |
| *mutability*.csv<br>*substitution*.csv<br>H (human)<br>M ( mouse) | Reference > Substitution model | Built-in | mutability and substitution matrices in csv format (cf SHazaM) | model of BCR evolution |
| Sequence_Patterns.csv | Reference | Built-in but editable | csv prefilled, column content can be changed additional columns can be added but will not be taken into account | plotting codon selection scores. Colors correspond to BCR sequence amino acid properties and PTM. |

**Supplementary Table 3.** Description of datasets used in this study.

| Source publication | Accession numbers | Technology | Species | tissue | #donors | Description | Associated figures | Short name |
| --- | --- | --- | --- | --- | --- | --- | --- | --- |
| Binet et al 2024<br>DOI:<br>[10.1101/2024.07.26.605240](https://doi.org/10.1101/2024.07.26.605240) | GSE275122<br>GSE275124<br>GSE275123 | 10x 5' | mouse | LN | 3 | Mice were immunized with NP-KLH and sacrificed at day 16 after immunization. Umi data available for QC filters. GC or GC-derived B cells were sorted with the AID-Cre-YFP reporter. | Fig.2<br>Fig.3B, Fig.3D,<br>Fig.4A, Fig.4B,<br>Fig.5B, Fig.5C,<br>Supp.Fig.2,<br>Supp.Fig.3,<br>Supp.Fig.4A,<br>Supp.Fig.4B,<br>Supp.Fig.6,<br>Supp.Fig.9,<br>Supp.Fig.10A-B,<br>Supp.Fig.11,<br>Supp.Fig.12A | NP-KLH1 |
|  |  |  |  |  | 2 | Mice were immunised with NP-KLH and sacrificed at day 16. prePC and PCs were enriched by FACs prior to sequencing. GC or GC-derived B cells were sorted with the AID-Cre-YFP reporter. | Fig.5F | NP-KLH2 |
|  |  |  |  |  | 2/<br>Time point | Mice were immunised with NP-KLH and sacrificed at day 7, 16, or 21. Umi data available for QC filters. GC or GC-derived B cells were sorted with the AID-Cre-YFP reporter. | Fig.6A,<br>Fig.6B,<br>Fig.6C | NP-KLH3 |
| Sprumont et al. 2023<br>DOI:<br>[10.1016/j.cell.2023.10.022](https://doi.org/10.1016/j.cell.2023.10.022) | Upon request | Plate based | mouse | LN | 6 | Mice were infected with influenza and sacrificed at day 18. GC and GC-derived PCs with measurable affinity for HA were sorted. No GEX library. | Fig.3E, Fig.5D,<br>Fig.5E,<br>Supp.Fig.4D,<br>Supp.Fig.10C,<br>Supp.Fig.10D | Mouse Influenza |
| King et al 2021<br>DOI:<br>[10.1126/sciimmunol.abe6291](https://doi.org/10.1126/sciimmunol.abe6291) | E-MTAB-8999<br>E-MTAB-9003<br>E-MTAB-9005 | 10x 5' | human | Tonsil | 5 | scRNAseq was performed on pediatric tonsilectomies. No cell enrichment. | Fig.3C, Fig.4C,<br>Supp.Fig.3E,<br>Supp.Fig.4A,<br>Supp.Fig.4B,<br>Supp.Fig.7 | Human tonsil |
| Kim et al<br>2022<br>DOI:<br>[10.1038/<br>s41586-022-04527-1](https://doi.org/10.1038/s41586-022-04527-1) | PRJNA7316<br>10<br>PRJNA7412<br>67 | 10x 5' | human | LN/<br>Blood | 15 | Donors with confirmed SARS-Cov2 infection provided fine needle aspirates of draining axillary lymph nodes. GC B cells, memory and PC were sorted. Elisa was performed on LN and blood derived monoclonal antibodies, with recombinant SARS-Cov2 spike protein. | Supp.Fig.8<br>Supp.Fig.14C | Human<br>covid |
| Chen et al<br>2022<br>DOI:<br>[10.1038/<br>s41586-020-2262-4](https://doi.org/10.1038/s41586-020-2262-4) | GSE140795 | Rep-SHM<br>Seq | mouse | PP | 10 | Mice repertoires were compared between WT and GF mice PP repertoires. Sequences from GF mice were used to compute the SHM targeting model. | Supp.Fig.5 |  |

**Supplementary Figure 1.**
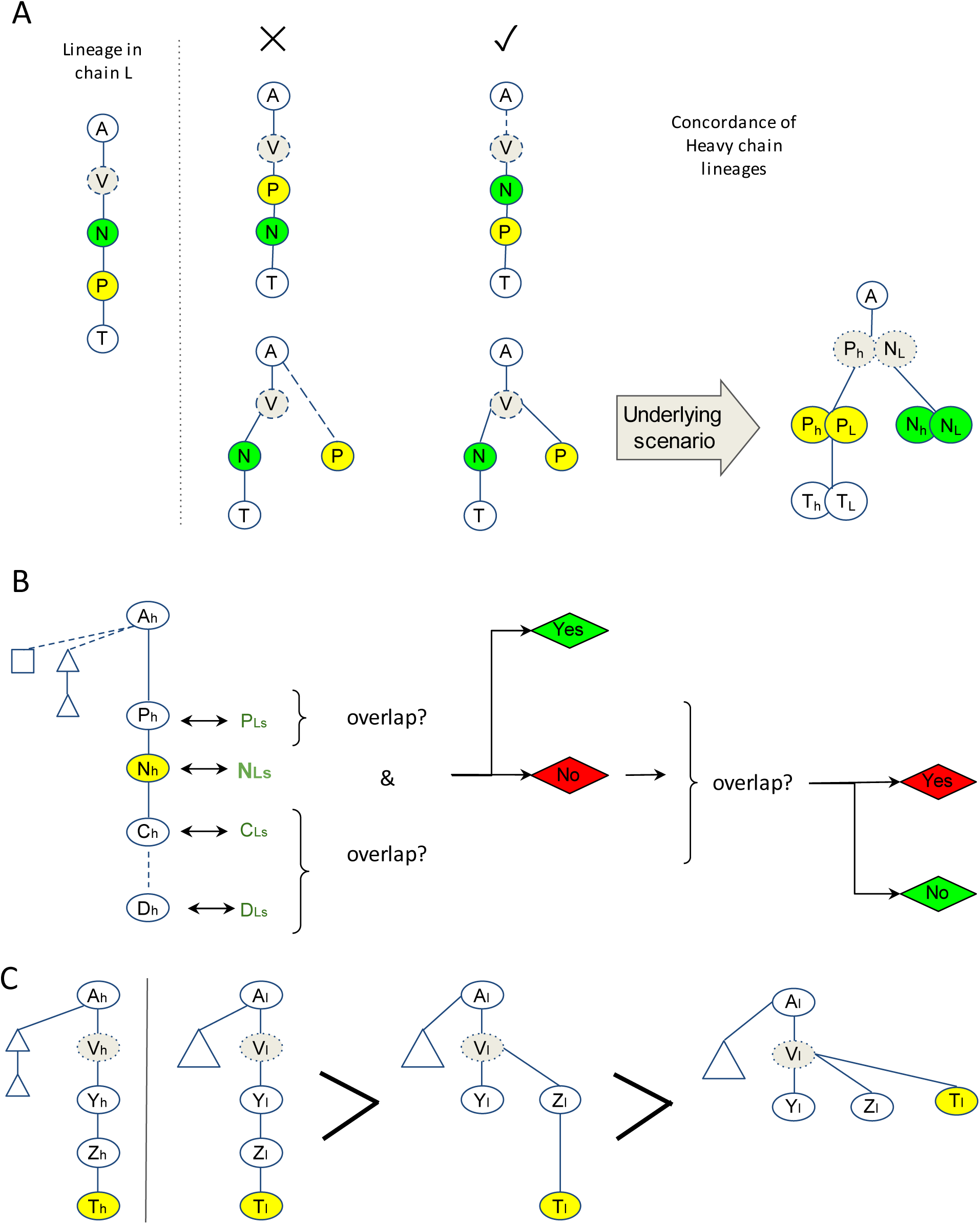
Details on tree selection implementation. **A)** Schematic representation of tree selection based on concordance criterion. SeQuoIA first checks whether BCR relationships are consistent in heavy chain and light chain trees. **B)** The order of paired light chains obtained from the heavy chain trees and the actual order in the light chain trees has to be conserved. If so, SeQuoIA checks that a paired node is not scattered in different sub-lineages in heavy and light chain trees, by checking that ascendance and descendance have common elements in both trees. If a sequence is both within the ascendance and descendance without a continuum, a penalty is applied to the pair of trees. **C)** Lineage overlap optimization among consistent tree pairs. P = Parent, N = node, C = Child, D = descendant, A = ancestor, T = terminal node, V = virtual node. H = heavy, L = light

**Supplementary Figure 2.**
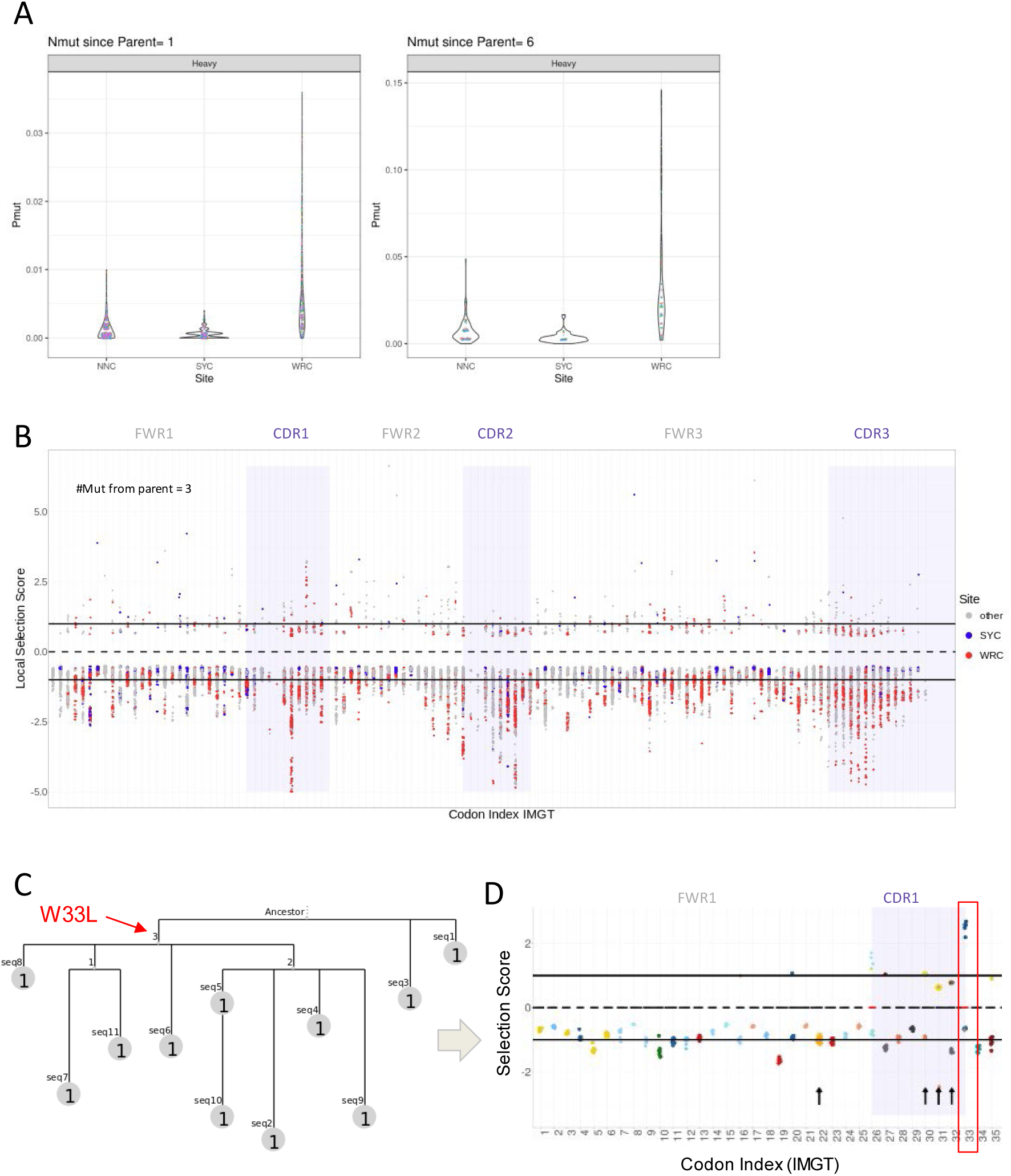
Local selection scores. All analyses are performed on mouse NP-KLH1 dataset **A)** Dotplot of raw mutation probabilities quality control of known hotspots and coldspots as provided by the SeQuoIA pipeline at the end of the mutation modeling step. **B)** Codon selection score distribution across heavy chain, all clonotypes combined. Solid lines materialize putative positive (upper part) and negative (lower part) selection. **C)** Phylogenetic tree of a clonotype featuring sequences with and without W33L mutation in its heavy chain. **D)** Local selection score distribution in the same clonotype. The acquisition of W (blue) mutation is associated with positive selection score whereas its maintenance does not appear to be the result of negative selection.

**Supplementary Figure 3.**
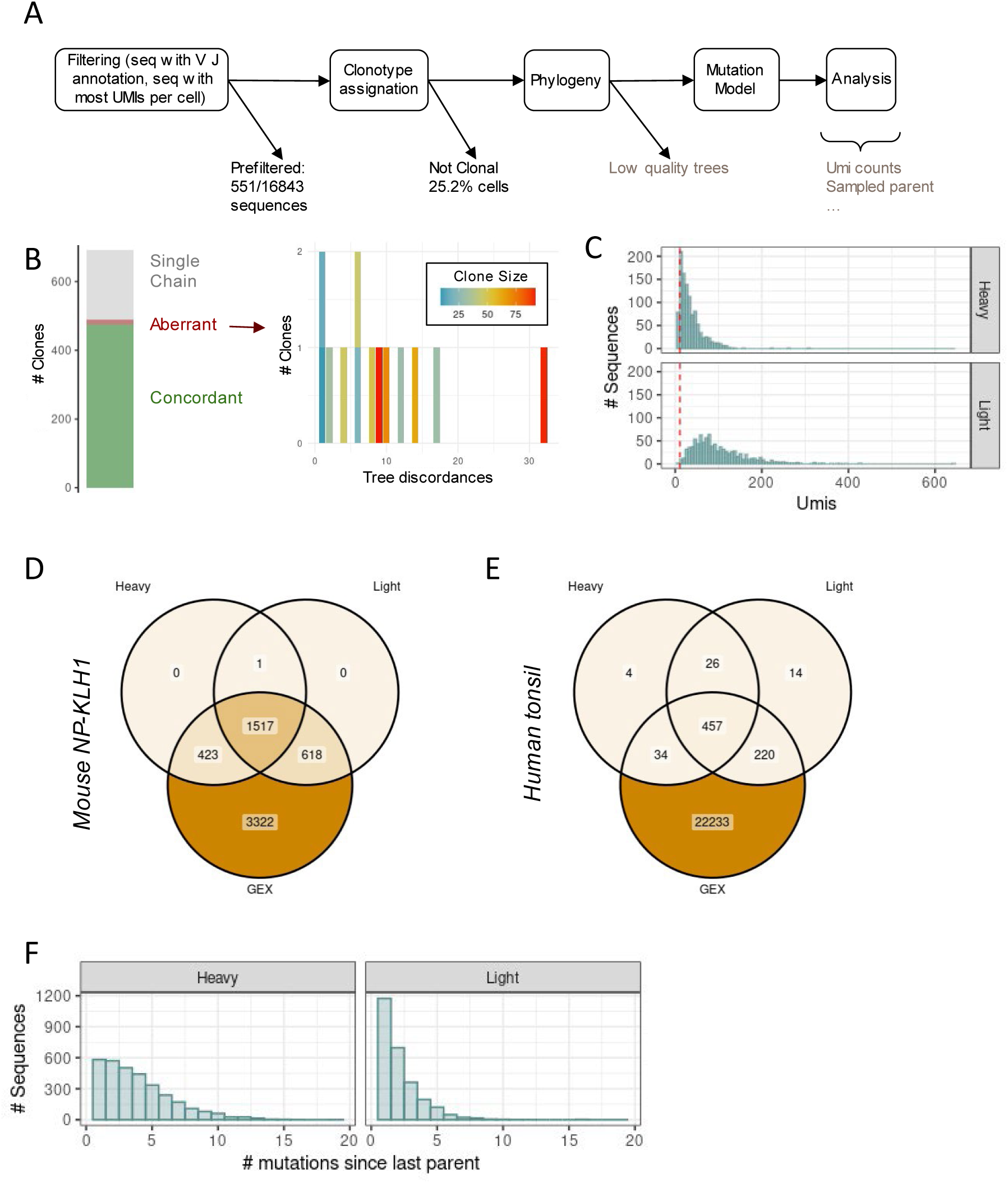
Features of integrative single cell datasets. **A)** BCR data loss after quality controls. Filtering steps in black are automated whereas filtering steps in gray are part of user’s choice. **B)** Quality of tree pairs within clonotypes. Right panel depicts clone size and the number of discordances. Clonotypes with concordant tree pairs were kept for following analyses. Trees with low discordance to clonotype size ratio were not discarded to compromise between data quality and data loss. **C)** BCR library UMI counts distribution for non-PC and non-proliferative cells. Cells with low UMI counts (< 15) were discarded. **D) E)** Single-cell gene expression (GEX) and BCR information availability in mouse (D) and human (E) datasets after filtering steps. **F)** Mutation from last parent in tree distribution. Low mutation counts is associated with enhanced modeling accuracy and is informative on instantaneous selection. All figures correspond to mouse NP-KLH1 dataset, unless stated otherwise.

**Supplementary Figure 4.**
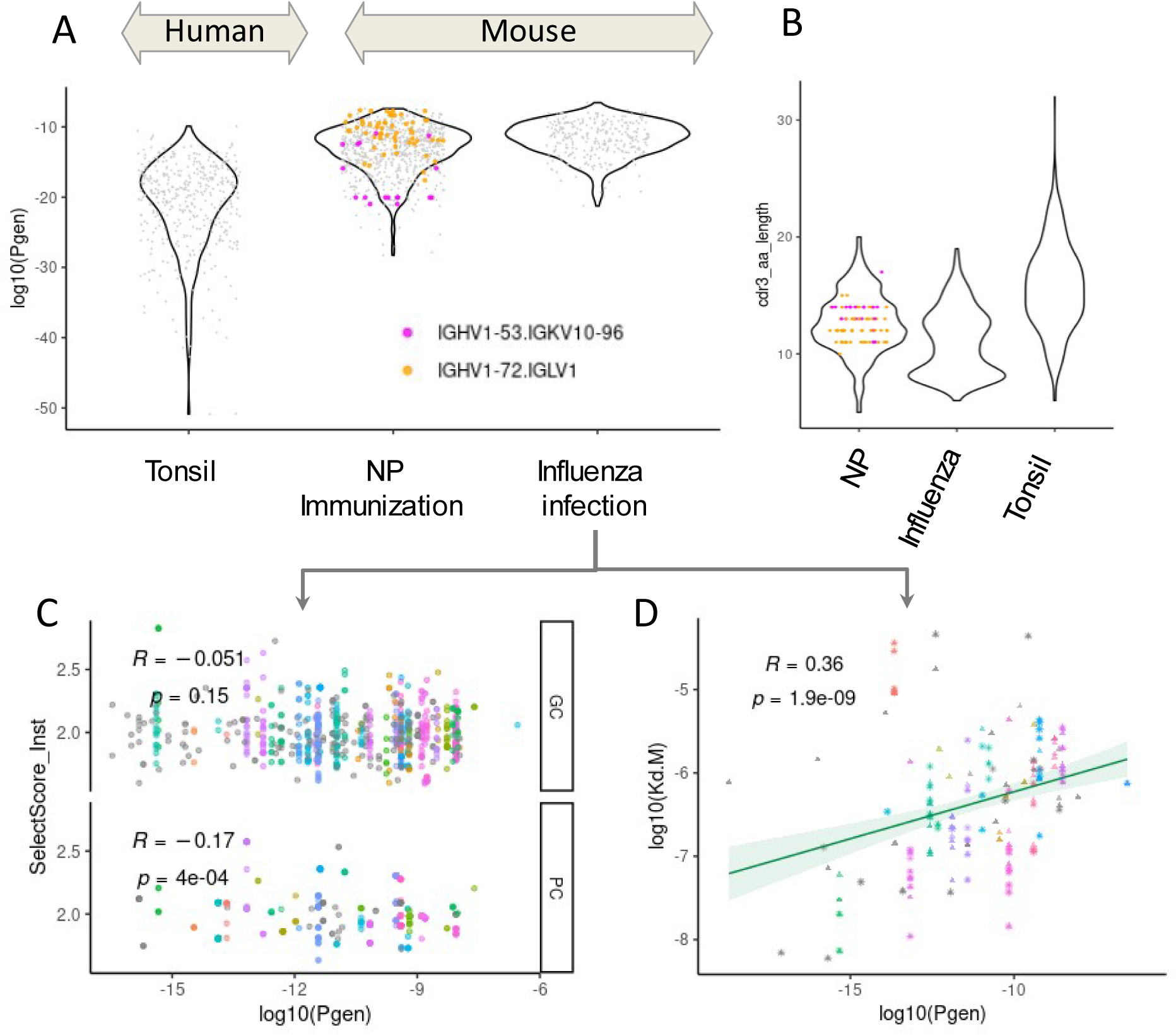
Application of IGoR within the SeQuoIA pipeline **A)** Pgen distribution of NCAs across datasets. Clonotypes recurrent in NP immunization model are highlighted. **B)** CDR3 length (nucleotides) comparison between datasets. **C)-D)** Spearman correlation of Pgen and affinity. Dots correspond to sequences with measured affinities and colors to clonotypes (within which all sequences have the same Pgen) in Sprumont et al. dataset. We compared the intrinsic generation probabilities (Pgen) of human and murine clonotypes to investigate whether chimeric clones were due to low of preGC selection. Murine clonotypes from both pauciclonal and polyclonal responses were associated with higher Pgen. Reasons for this could be either a mouse-specific intrinsic bias in VDJ recombination, or permissive pre-GC selection where rare clones would not be especially favoured. In murine models, competition for follicular niches and antibody feedback should be limited as there is no infection history. In line with the first hypothesis, CDR3 lengths were smaller in mice than humans, thus limiting junctional diversity. IGOr Pgen inference was validated in figure C, where Pgen are inversely correlated to the affinity, suggesting that most probable combinations are not present due to a selection process.

**Supplementary Figure 5.**
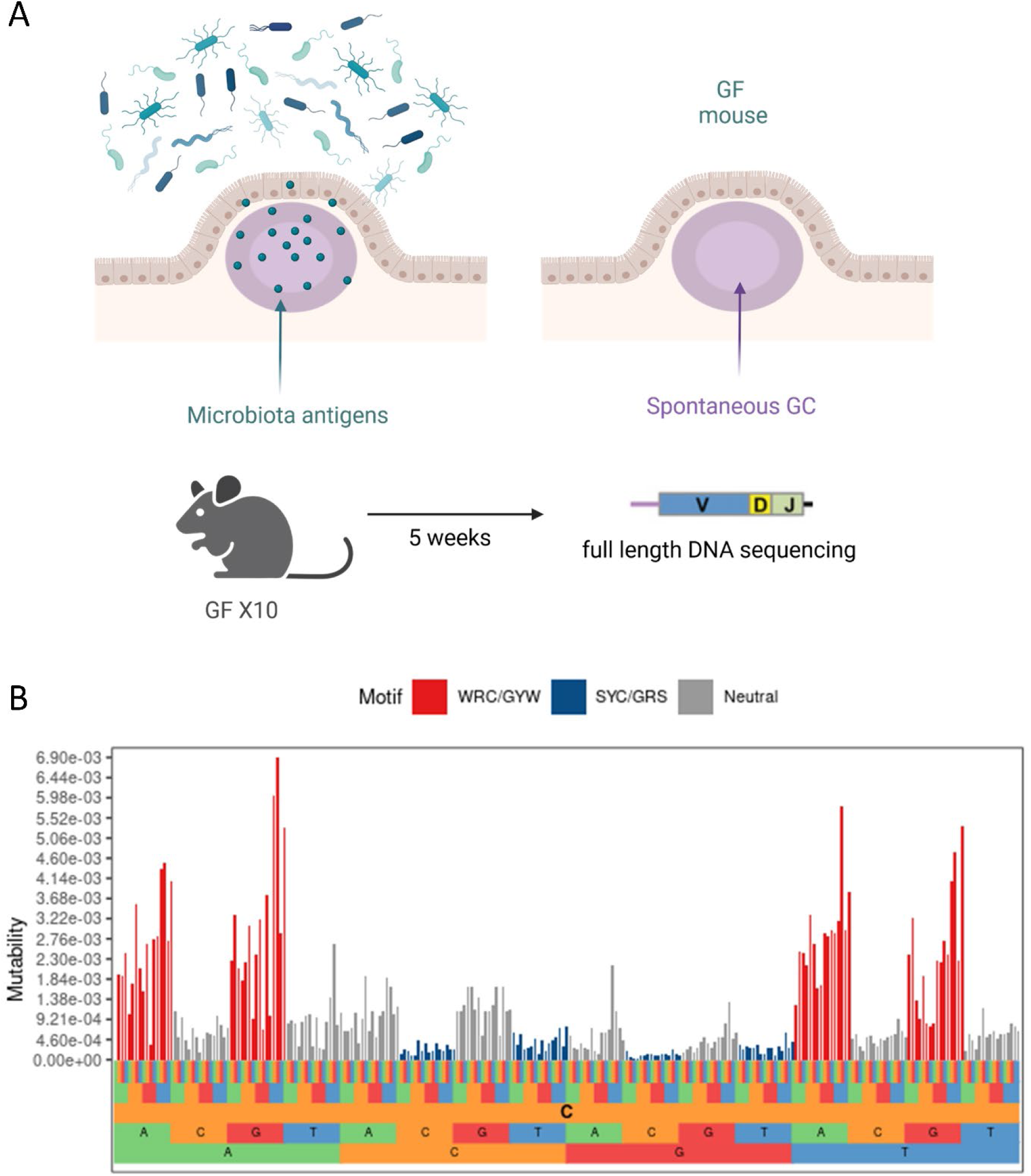
Construction of a new murine AID mutation model. **A)** Schematic representation of scBCRseq experiment in PP dataset from Chen *et al.* Only Germ-free mice (GF) sequences were kept to build the mutation model since microbiota antigen binding does not confer a selective advantage to Peyer’s patches GC B cells. **B)** SHM targeting patterns inferred with ShazaM tool. Letters in the lower parts correspond to 5-mer context of targeted nucleotide. Well described mutation hotspots and coldspots are depicted in red and blue respectively, as a quality control.

**Supplementary Figure 6.**
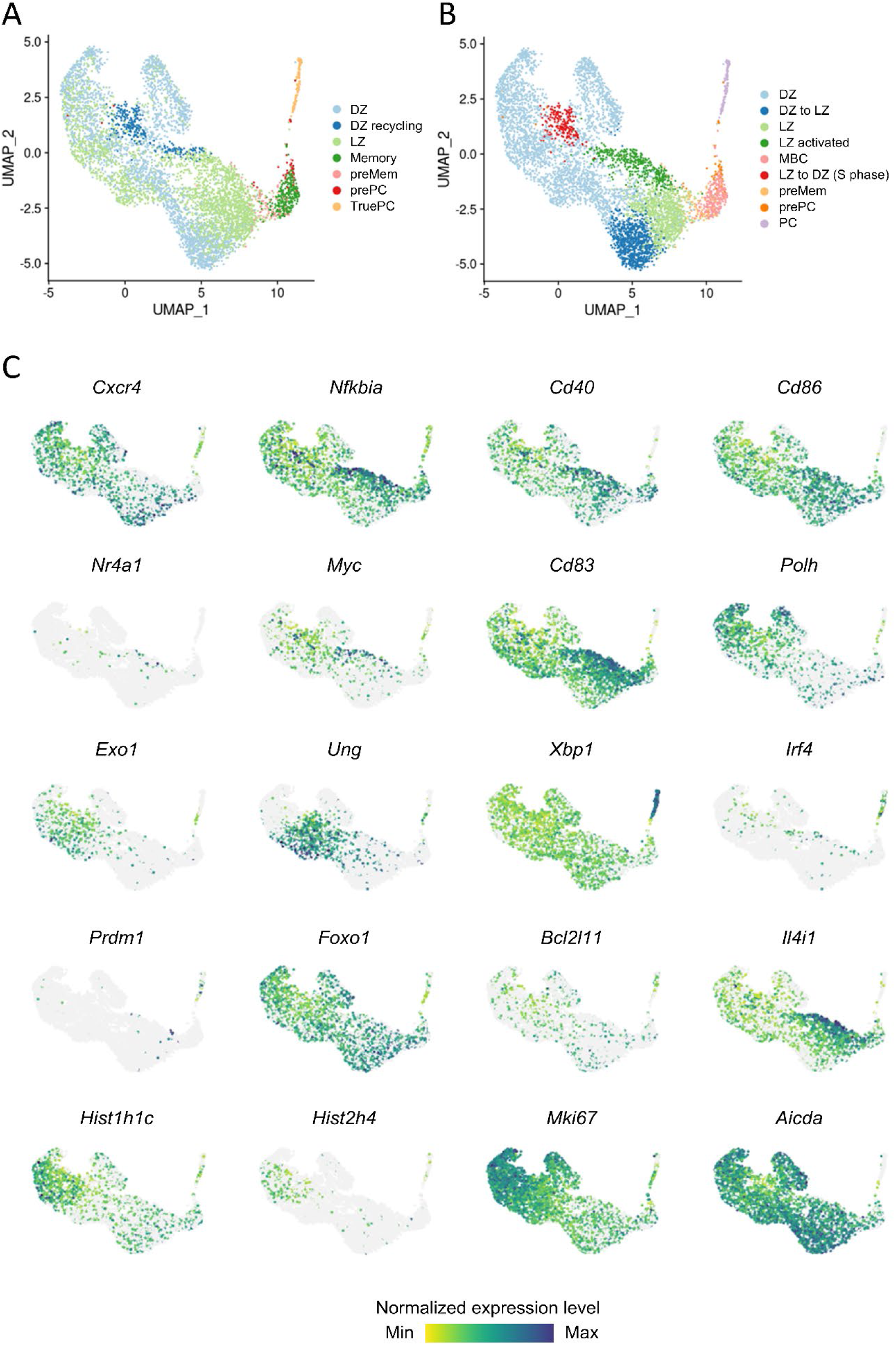
scRNA-seq annotation of NP-KLH1 dataset. **A)** UMAP projection of scRNAseq data colored by original annotations from Binet et *al*. **B)** and by our cluster-based annotation. **C)** Feature plots of marker genes. Gray dots correspond to cells with zero counts for the considered gene.

**Supplementary Figure 7.**
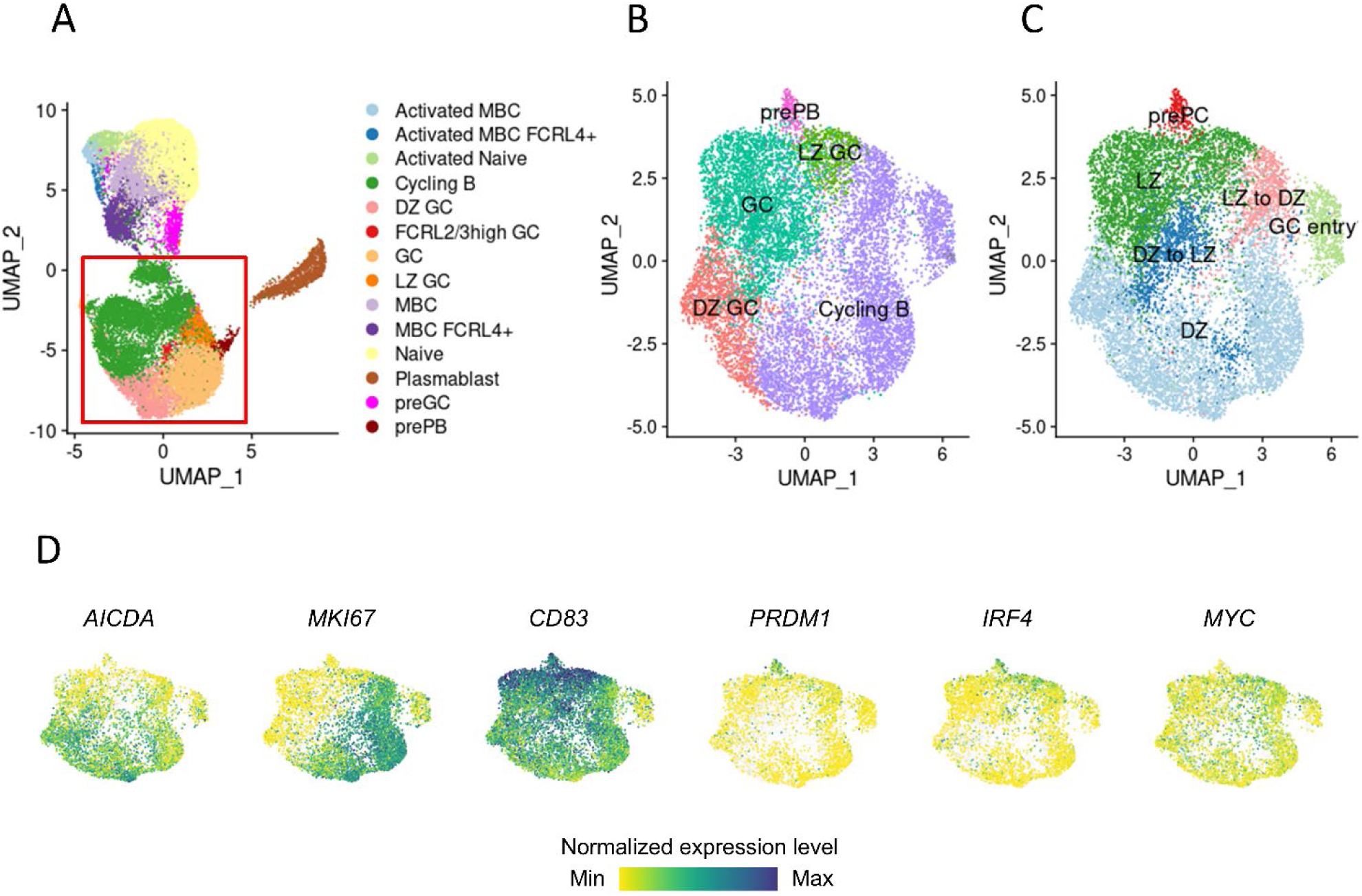
scRNAseq annotation of human tonsil dataset. **A)** UMAP projection of tonsillar B cells after cell cycle regression. Cell color corresponds to original annotation from King et *al*. Cycling and GC B cells were subsetted (red square) for further annotation **B)** UMAP projection of subsetted cells with original annotations. **C)** Same as B) with cluster-based reannotations. **D)** Feature plots of marker genes expression.

**Supplementary Figure 8.**
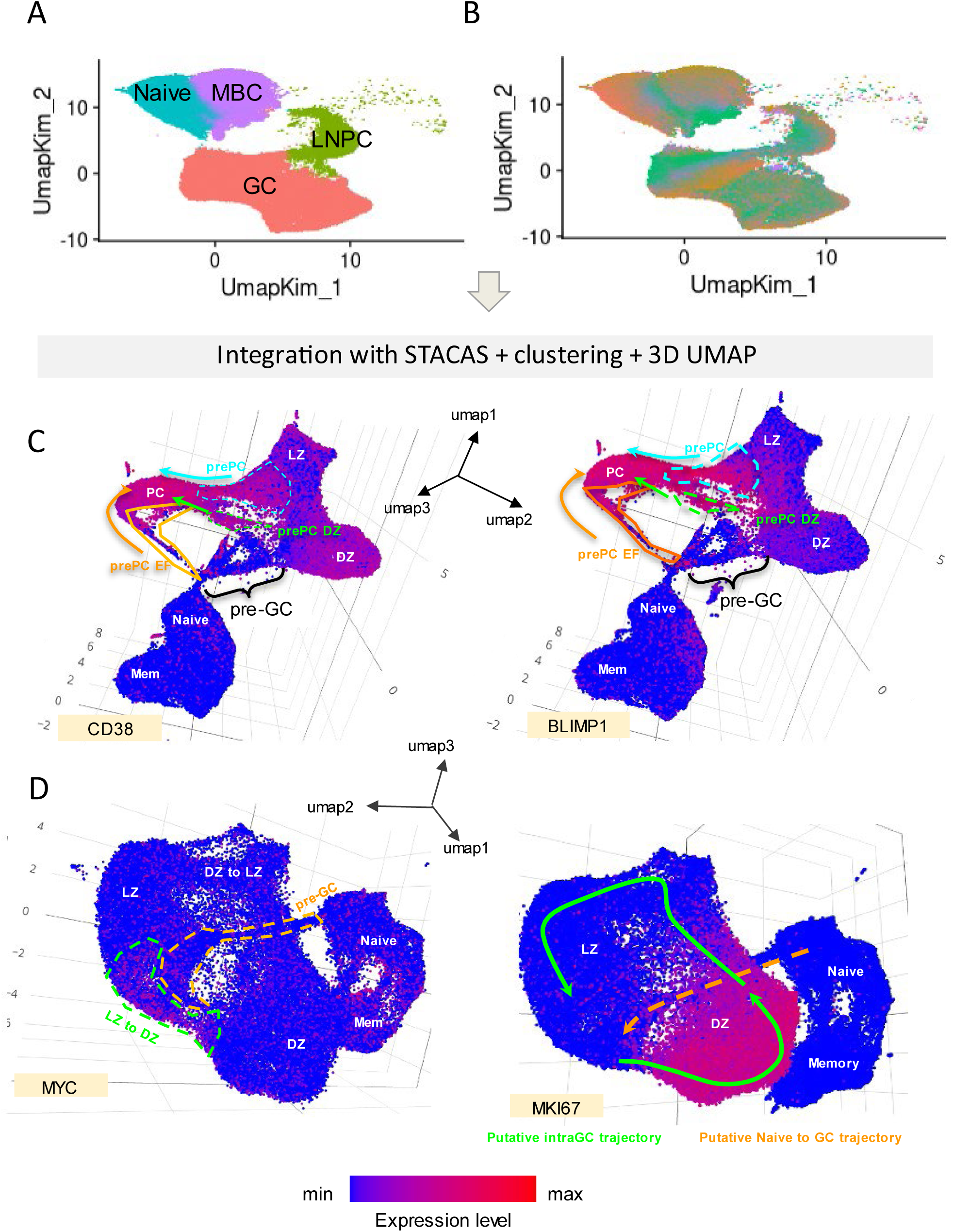
scRNAseq annotation of human Covid dataset. **A)** original UMAP projection and annotations of scRNAseq data from Kim et *al*. **B)** Original UMAP projection coloured by donor. **C)** prePC annotation. Three intermediary clusters bridging PCs to other clusters are highlighted in the 3D UMAP. CD38 expression levels support GC-experienced cells identification. BLIMP1 expression supports PC and pre-PC identification. **D)** LZ to DZ cluster annotation and heterogeneity. NB: putative trajectories were conjectured based on current knowledge of EF and GC dynamics.

**Supplementary Figure 9.**
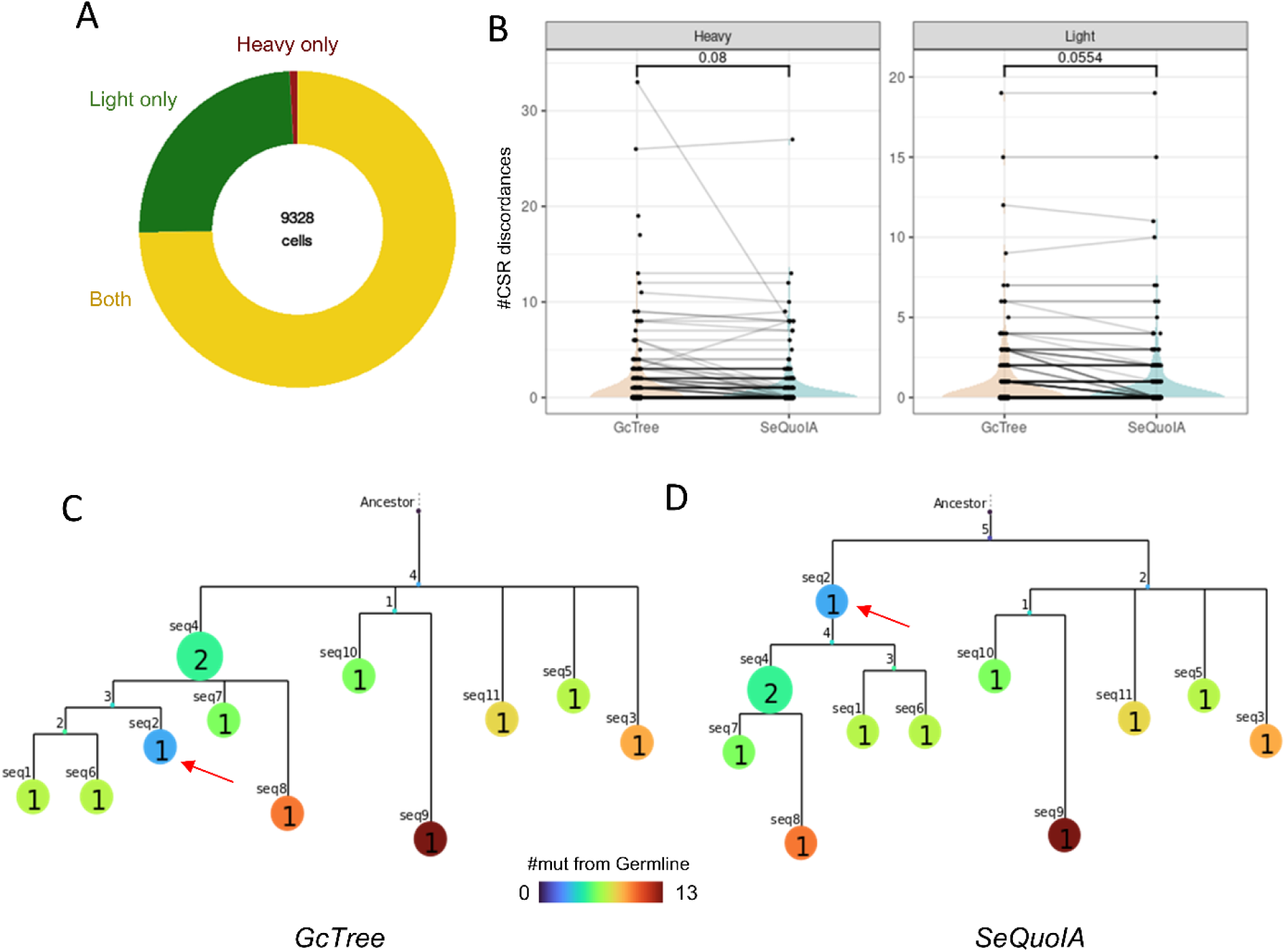
Complementary to Figure 2. **A)** Chain availability at the single cell level in mouse NP-KLH1 dataset. **B)** Optimization of CSR order in phylogenetic tree selection. P values denote paired Wilcoxon rank sum test. **C)** Mutation load of BCR sequences in a tree selected with GcTree. The red arrow highlights an aberration in mutational load. **D)** Same as C) with the tree selected by SeQuoIA for the same clonotype. The aberration (red arrow) has been corrected with SeQuoIA.

**Supplementary Figure 10.**
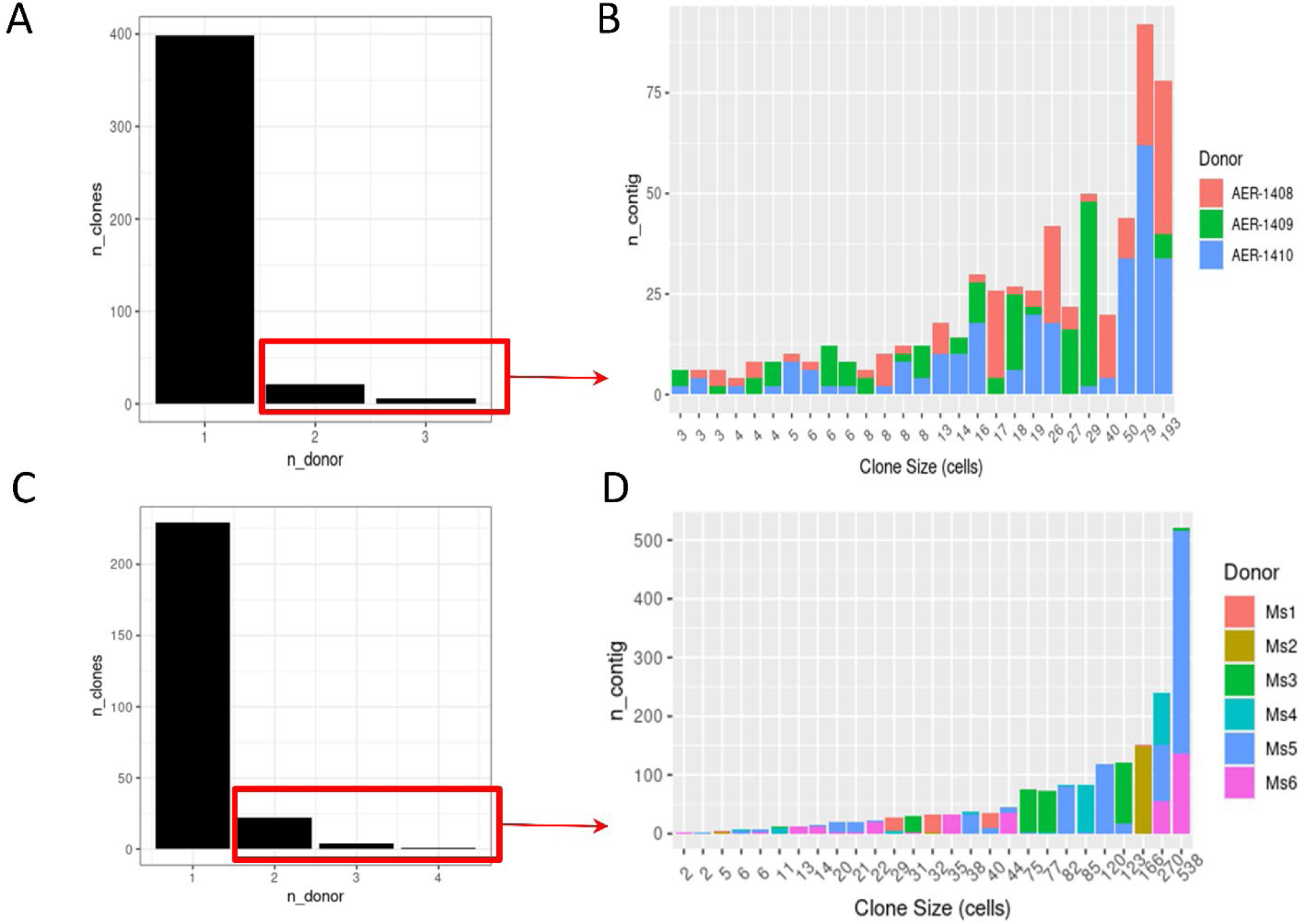
Clonotype assignation errors and optimization **A)** Number of clones featuring sequences from 1 or more donors. Clonotype assignation was performed on full BCR dataset with empirical 80% CDR3 similarity threshold in a pauciclonal dataset (mouse NP-KLH1). **B)** Barplot of sequence origin in clonotypes shared among several donors. **C) D)** Same as A-B in a polyclonal mouse influenza dataset.

**Supplementary Figure 11.**
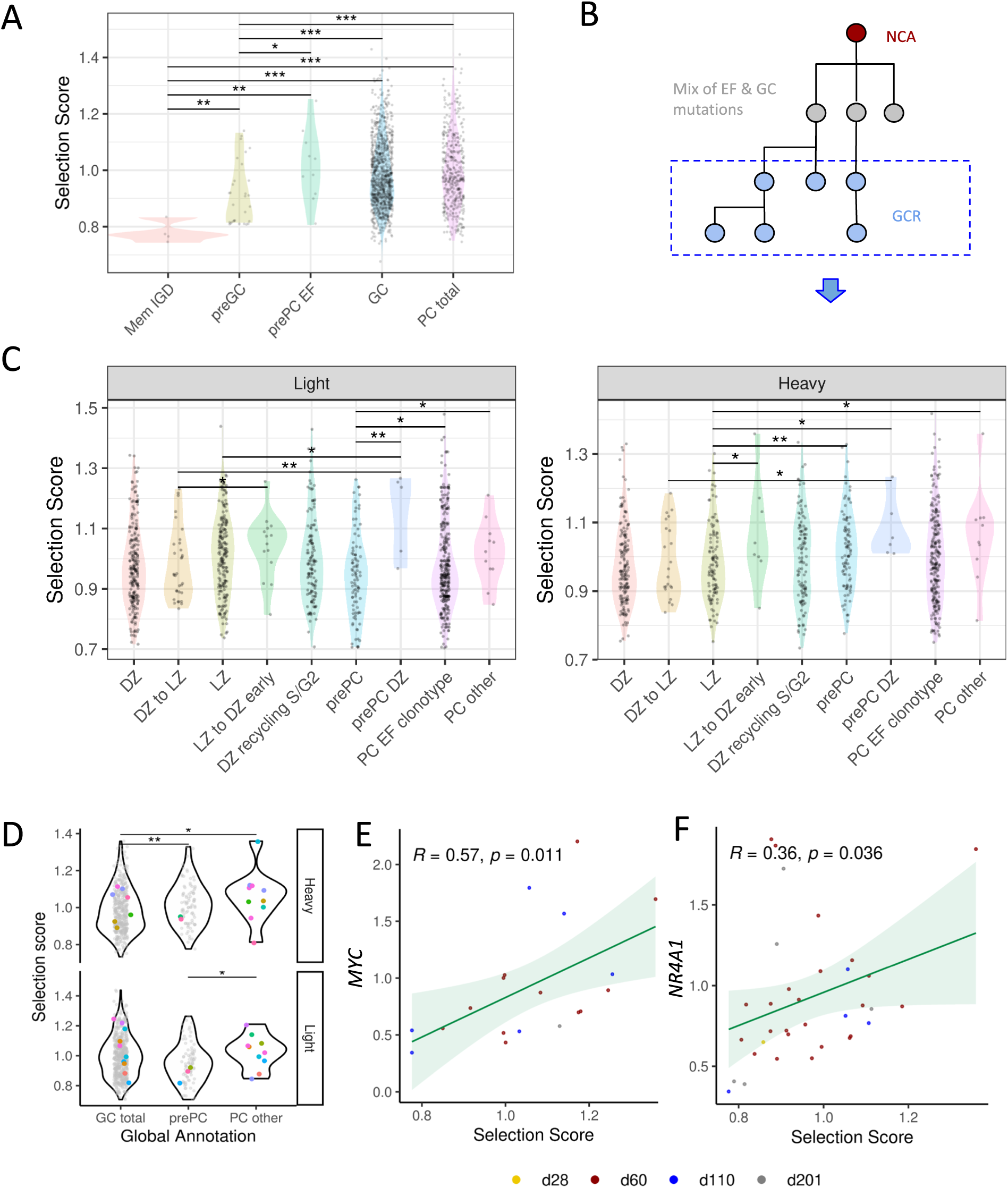
Application of SeQuoIA in human Covid dataset **A)** Heavy chain selection score distribution over general B cell subsets in LNs **B)** Schematic of tree-based enrichment of B cell undergoing GCR among total mutated B cells in human LNs. This strategy allows notably to discard Myc+CD38-B cell entering GCs clustering with LZ to DZ Myc+CD38+ cells **C)** Heavy and light chain selection score distribution across scRNAseq GC B cells clusters. NB: d60 time point was selected for this analysis as it makes up the major part of the whole dataset. Clonotypes with more than 5 aberrations in phylogenetic trees were excluded. PCs sharing the same clonotypes as EF prePCs were annotated as a separate cell subset. Remaining PCs should thus be enriched in GC-derived cells. Each dot represent a BCR of a given phenotype 1 or 2 mutations away from its parent. **D)** Unique BCR sequence selection score (heavy and light chains) comparison between putative GC-derived PCs and total GCB cells. Clonotypes present in PC compartment are highlighted with different colors. **E)F)** Spearman correlation between gene expression level and heavy chain selection score. All p-values denote wilcoxon rank-sum tests. All timepoints were taken for this analysis and indicated with different colours.

**Supplementary Figure 12.**
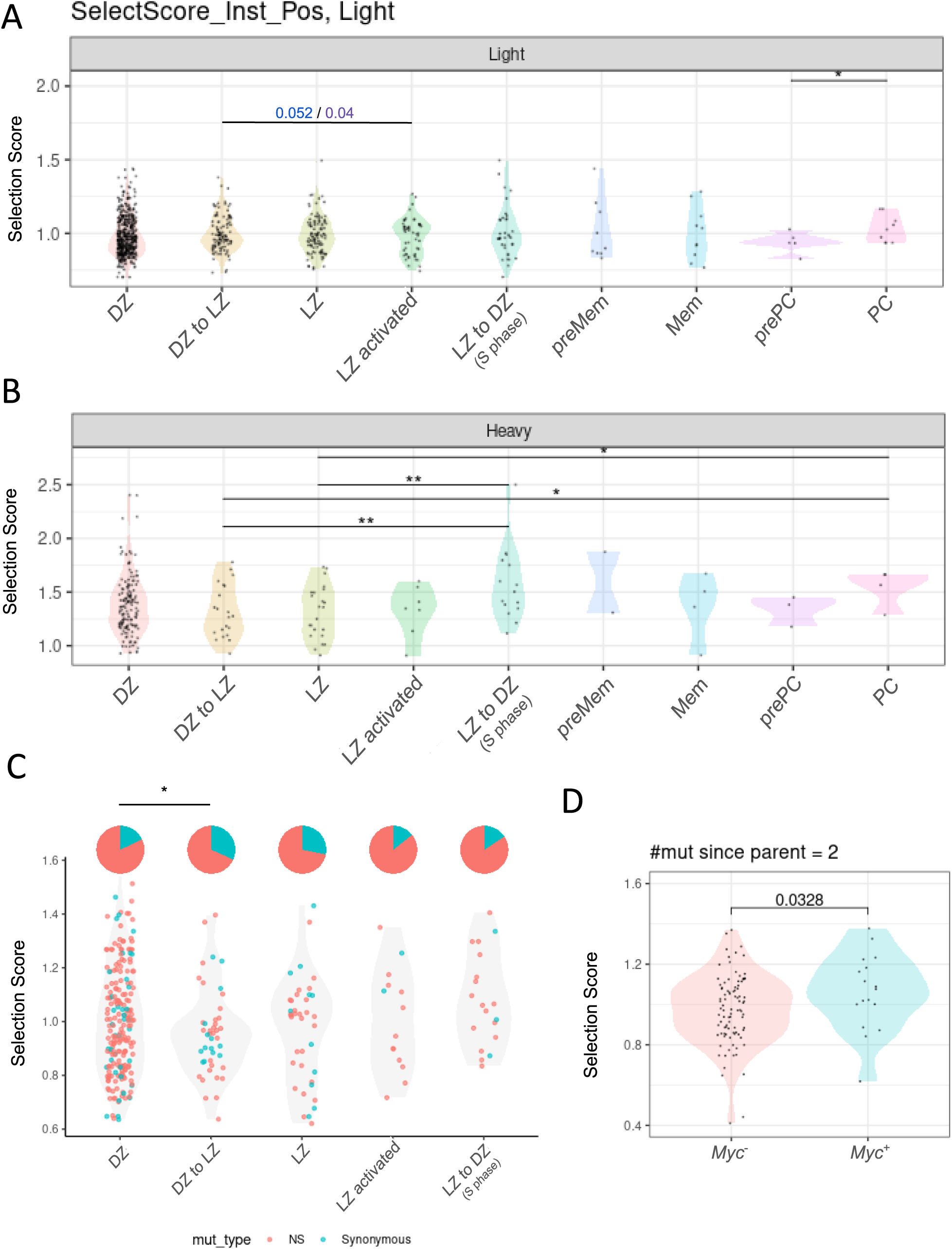
Complementary to Figure 3 **A)** Light Chain selection score distribution across LN B cell subsets. p-values in blue correspond to Kolmogorov-Smirnov distribution comparison tests and p-values in purple correspond to a proportion (of scores >1) test. **B)** Same as fig. 3.B considering only non synonymous mutations. **C)** Synonymous and NS mutation ratios and heavy chain selections scores across subsets. P-value corresponds to one-sided binomial test on proportions (of NS mutations). **D)** Heavy chain selection scores comparison between *Myc*+ and *Myc*-cells in the LZ. p-values in black denote wilcoxon rank sum test on means. All figures correspond to mouse NP-KLH1 dataset.

**Supplementary Figure 13.**
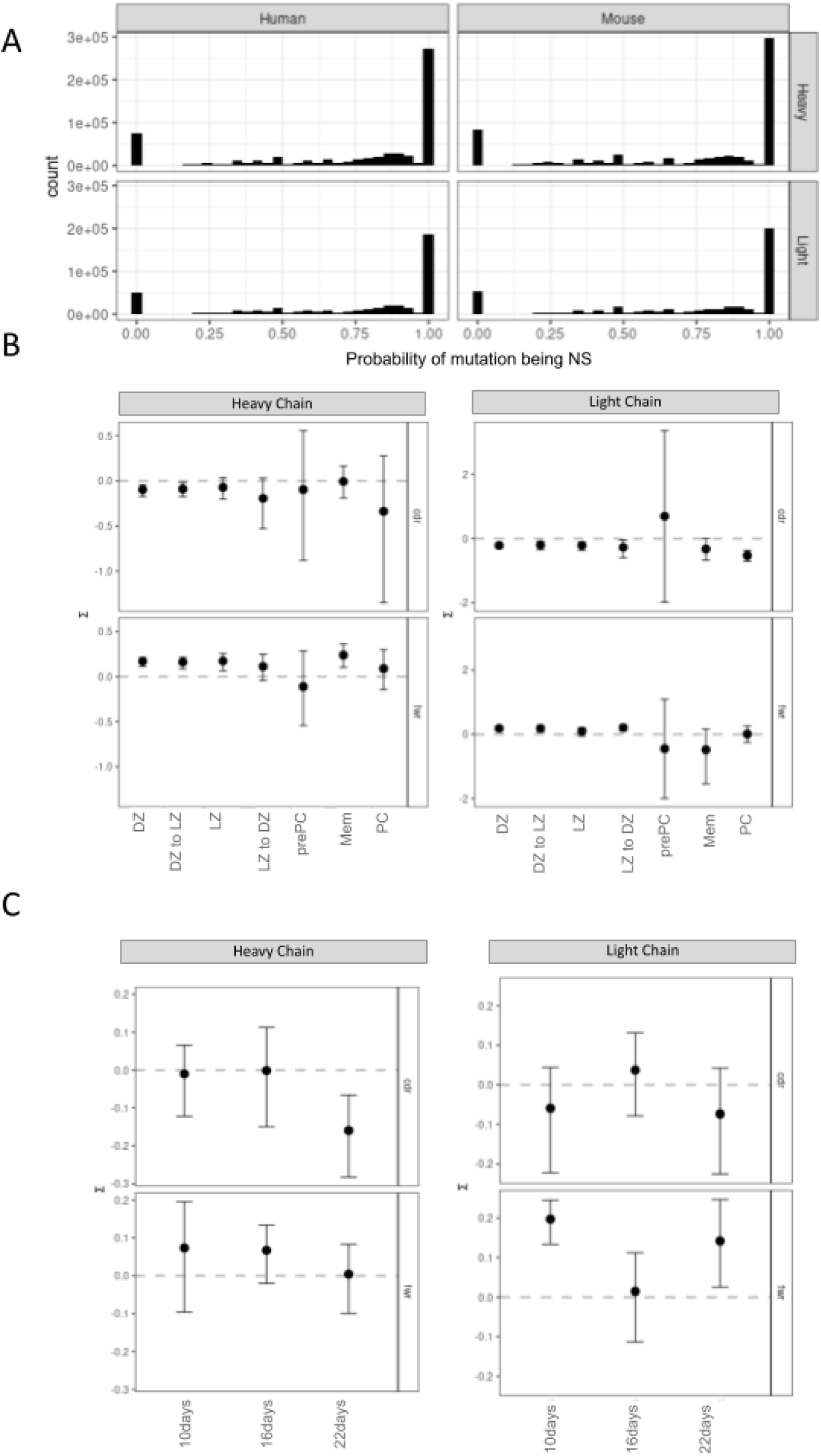
Performance of BASELINe. **A)** NS mutation probabilities distribution across codon sites in the absence of selection in human tonsil (left) and mouse NP-KLH1 (right) datasets **B)** BASELINe selection quantification across B cell subsets in mouse NP-KLH1 dataset. **C)** BASELINe selection quantification across post-immunization time points in mouse NP-KLH3 dataset.

**Supplementary Figure 14.**
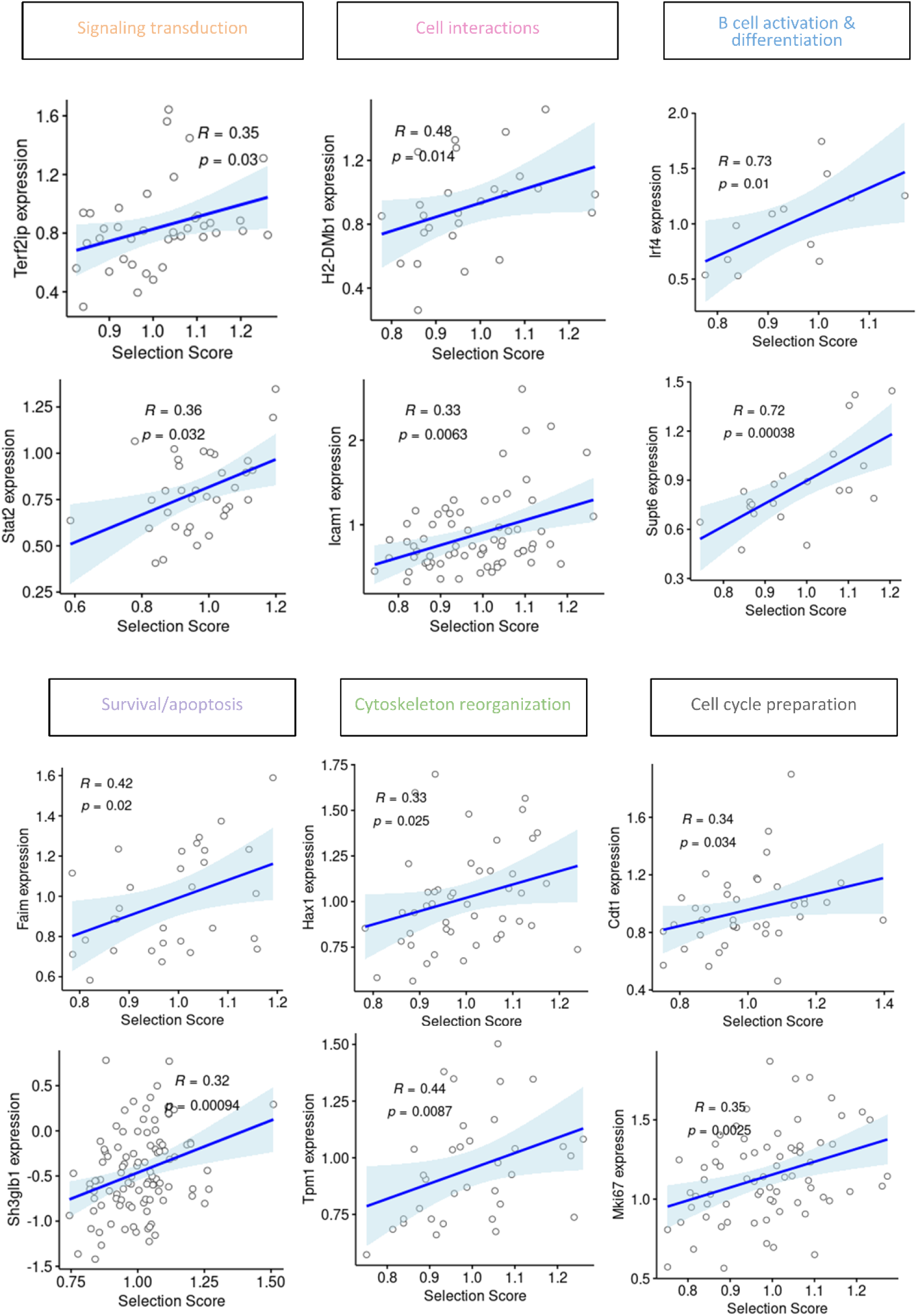
Candidate genes that could provide a selective advantage to GCB cells based on BCR properties. Spearman correlation between light chain BCR selection score (1 mut away from parent) and LZ gene expression from GO pathways from Figure 4.

**Supplementary Figure 15.**
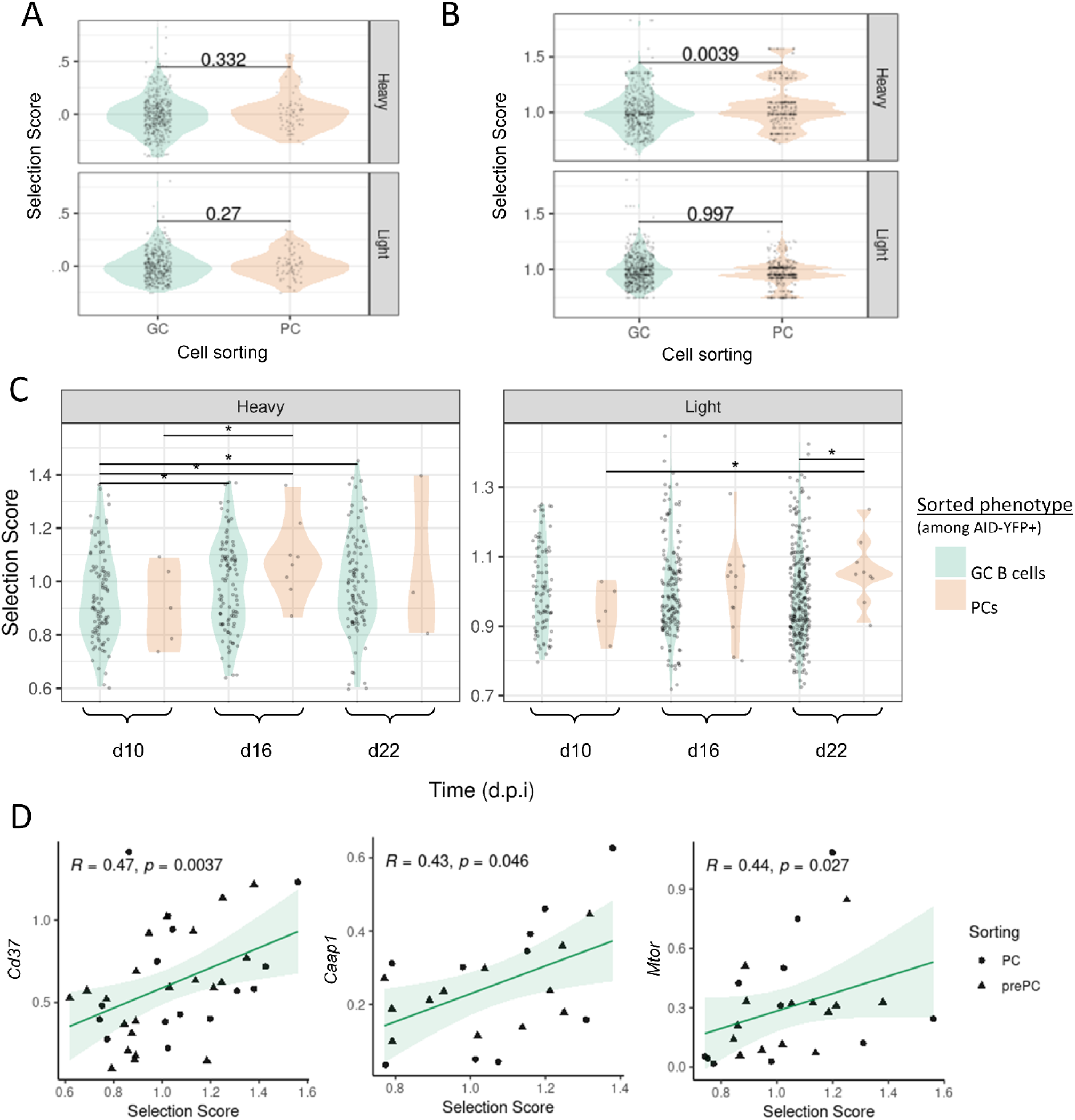
PC selection study across datasets. **A)** PC and GC B selection scores in mouse influenza dataset. Unique sequences are taken. P-values denote wilcoxon ranked sum test. **B)** Same as A) with total BCR sequences so as to consider proliferative effect **C)** PC and GC B cell selections scores distribution at different timepoints in NP-KLH3 dataset. Unique Ig sequences were taken. P-values denote wilcoxon rank sum tests (uncorrected) **D)** Spearman correlation between prePCs/PCs selection scores and selected genes in NP-KLH3 dataset (enriched for these two cell types). NB: score correlation was extended to 2 mutations since parent for Caap1

**Supplementary Figure 16.**
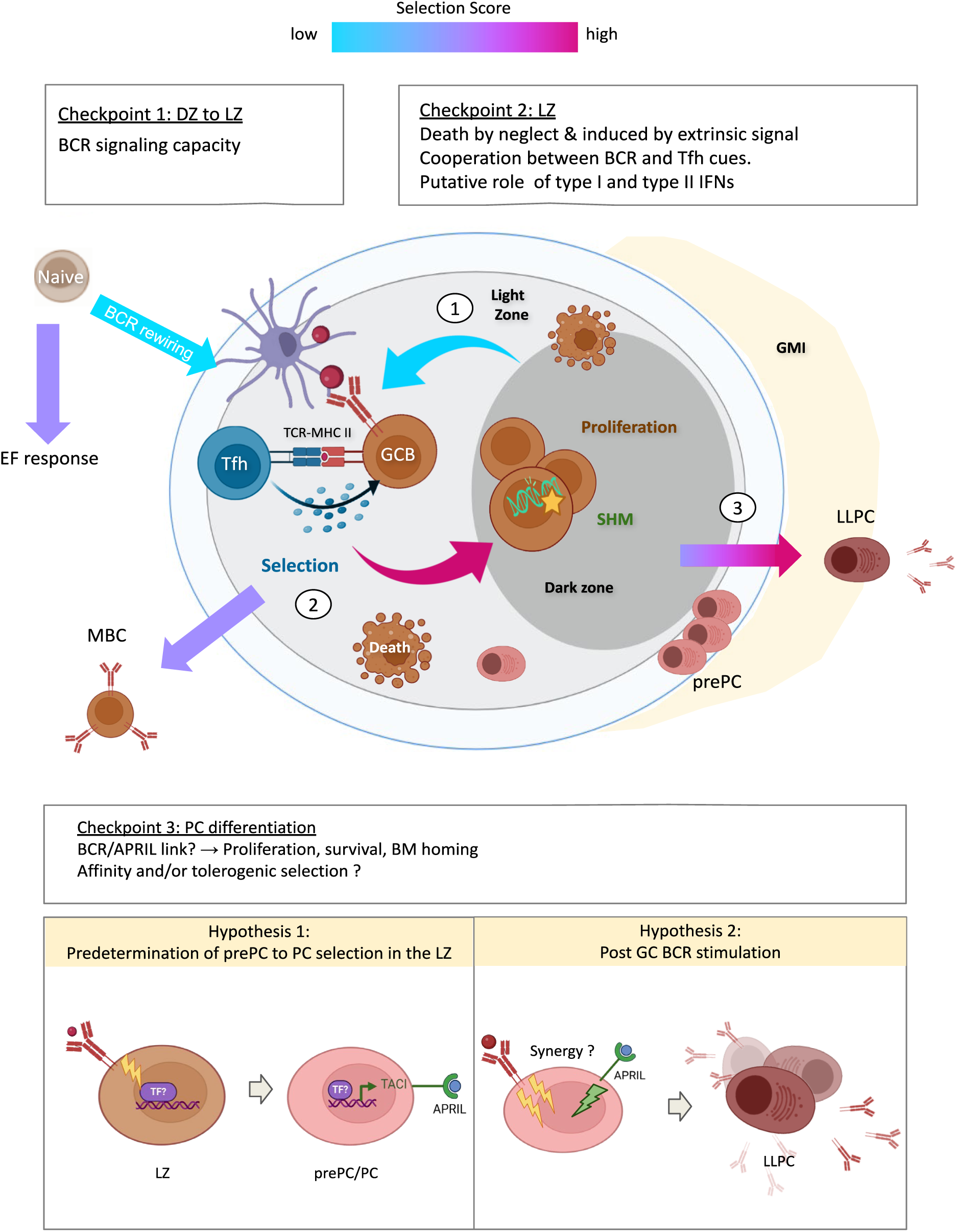
Working model of GC or GC-derived B cell selection. Circled numbers correspond to successive selection bottlenecks identified with selection score distribution across scRNAseq clusters. GMI = GC/medulla interface. LLPC = long lived plasma cell.

